# An Integrated Preclinical Platform for Lethal Neuroendocrine Prostate Cancer from Rapid Autopsy Bone and Liver Metastases

**DOI:** 10.64898/2026.07.22.740121

**Authors:** Borum Ryu, Thomas C Caffrey, Sriya Sridhar, Catherine S Johnson, Ramia J Salloom, Kabhilan Mohan, Grace Waldron, Amanda Robotham, Ethan M Wilcox, Diane Costanzo-Garvey, Janice Taylor, James Talaska, Rachel Rhatigan, Quan P Ly, Heather C Smith, Kaustubh Datta, Surinder K Batra, Chad A LaGrange, Benjamin A Teply, Subodh M Lele, Michael A Hollingsworth, R Katherine Hyde, Kyle J Hewitt, Gargi Ghosal, Fanben Meng, Angie Rizzino, Adrian R Black, Paul M Grandgenett, Maher Y Abdalla, Leah M Cook, Raymond C Bergan, Grinu Mathew

## Abstract

Treatment-emergent neuroendocrine prostate cancer (NEPC) is an aggressive, therapy-resistant disease arising in up to 20% of castration resistant prostate cancers, yet robust biologically relevant preclinical models remain scarce. Here, we describe a technical blueprint for establishing an integrated platform of patient-derived models from visceral and bone metastases collected through a prostate cancer rapid autopsy program (PC RAP). We report the establishment and characterization of patient-derived xenograft (PDX) models from liver metastasis tissue, liver and bone metastasis-derived organoid lines (PDOs), and corresponding patient-derived organoid xenograft (PDOX) models. In addition, we established, to our knowledge, the first mesenchymal stem cell (MSC) cultures derived from neuroendocrine prostate cancer (NEPC) bone metastases. The PDOs preserved intratumoral heterogeneity, displaying both CRPC-NE and CRPC-adenocarcinoma features. These organoids retained neuroendocrine identity across multiple passages, with transcriptomic profiles concordant with the original patient tissue and matched PDX models generated at our institution and at the National Cancer Institute (NCI Patient-Derived Models Repository). To model the bone metastatic microenvironment, we generated novel organoid-based New Approach Methodologies (NAMs) by co-culturing PDOs with iPSC-derived bone marrow organoids, establishing a physiologically relevant vascularized organotypic model of PC bone metastasis. To extend our studies in vivo, we established preclinical models using the liver and bone metastasis-derived organoid models. The PDOX models were tumorigenic and developed spontaneous lymph node metastases, providing clinically relevant models for investigating lethal NEPC biology. Together, these complementary patient-derived models provide a robust and versatile platform for investigating NEPC biology, metastatic progression, and evaluating new therapeutic strategies.

**Graphical abstract:** 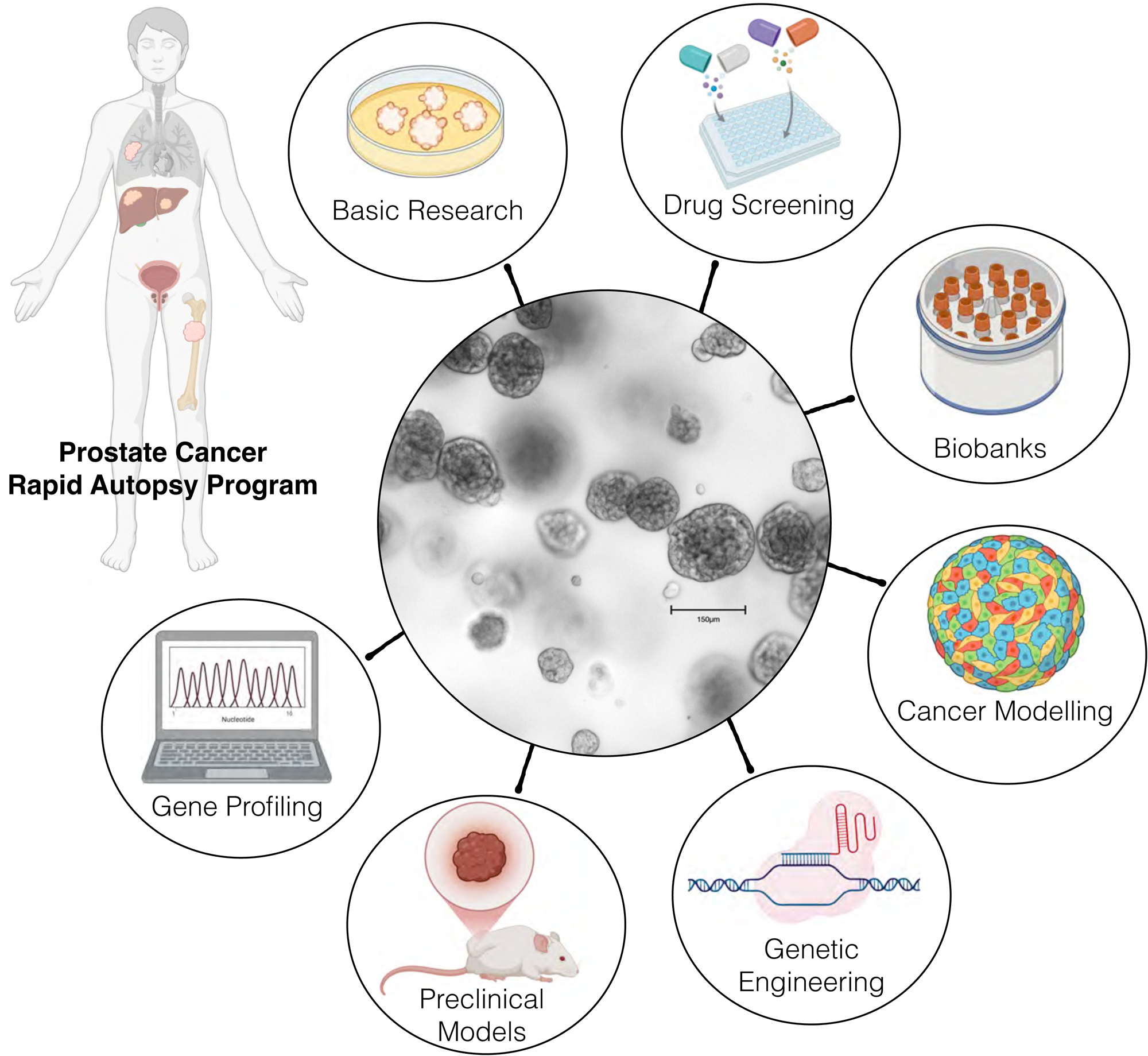

*Highlights:*

- Novel preclinical models of visceral and bone metastases established from a prostate cancer rapid autopsy program.
- This study is the first to establish mesenchymal stem cell cultures from NEPC bone metastases.
- PDOs preserve heterogeneity, showing both CRPC-NE and CRPC-Adeno features, with transcriptomic profiles concordant with originator tissue and PDX models.
- PC RAP-derived organoids are tumorigenic in vivo and generate spontaneous lymph node metastases.

## INTRODUCTION

Prostate cancer (PC) is the most commonly diagnosed malignancy in men and the second leading cause of cancer related death in the United States [1]. The vast majority of patients with PC initially present with androgen dependent adenocarcinoma, for which androgen deprivation therapy (ADT) remains the standard first-line treatment. While initial responses to ADT are nearly universal, castration resistance invariably develops through a series of adaptive mechanisms, most often involving reactivation or bypass of the androgen receptor (AR) signaling [2]. The widespread use of second-generation AR signaling inhibitors (ARSI) such as enzalutamide and abiraterone has dramatically reshaped the castration resistant prostate cancer (CRPC) landscape. However, this therapeutic pressure has also accelerated the emergence of AR-independent disease states [3, 4]. Among these, neuroendocrine prostate cancer (NEPC) represents the most clinically aggressive and molecularly distinct subtype, arising in up to 10-20% of patients with CRPC as a mechanism of resistance to AR targeted therapy [5]. Treatment-emergent NEPC (CRPC-NE) evolves clonally from prostate adenocarcinoma during disease progression, retaining PC genomic alterations such as loss of tumor suppressors PTEN, TP53 and RB1 while acquiring distinct epigenetic and pathway level changes that collectively define a neuroendocrine cell identity [6]. Patients with NEPC show a poor prognosis, with a median survival of less than two years. Treatment options are mainly untargeted, reflecting the absence of approved targeted therapies for treatment-emergent PC [5, 7].

Progress in understanding NEPC biology and identifying new therapeutic targets has been substantially hindered by the lack of reliable and representative preclinical models. For decades, the NCI-H660 cell line derived from the cervical lymph node of a patient with small cell carcinoma has been the only commercially available NEPC-like cell line [8–11]. Although patient-derived xenograft (PDX) models such as those in the LuCaP cohort have provided important insights into the genomic and phenotypic diversity of advanced PC, their utility for mechanistic studies and high-throughput drug screening is limited by the logistical demands of in vivo propagation and the difficulty of adapting PDX derived cells to in vitro culture [12, 13]. The development of three-dimensional organoid culture systems has fundamentally advanced the field. Organoids capture the spatial architecture, molecular identity, and, to some extent, the heterogeneity of the source tissue in a renewable, experimentally amenable format [14, 15]. Originally pioneered for intestinal epithelium using Lgr5 positive stem cells [16], organoid methods were progressively adapted for a range of human epithelial tumors, including colorectal [17], pancreatic [18], breast [19], and liver cancers [20], and eventually extended to PC [21, 22]. Establishing patient-derived organoids from metastatic CRPC has remained particularly challenging. Success rates from needle biopsies are modest, approximately 10-16% for fully characterized, long-term lines. This is due to limited input material, the selective pressures of culture conditions, and the tendency of rapidly dividing non-tumor cells to outcompete cancer cells during early passages [23, 24]. Optimization of culture conditions, for e.g. removal of the p38 MAPK inhibitor SB202190 and the antioxidant N-acetylcysteine (NAC) from prostate organoid media, has improved the establishment rate of CRPC cell lines derived from both patient biopsy, tissue, and PDXs [13]. The resulting organoid biobanks have demonstrated preservation of genomic copy number alterations, somatic mutation profiles, transcriptomic identity, and lineage marker expression relative to the originating PDX or patient tumor [13, 24]. The development of NEPC specific organoid models has demonstrated the power of this approach for rare and poorly understood cancer phenotypes. Early patient-derived NEPC organoids were developed from needle biopsies of metastatic lesions from four patients; molecular characterization was possible in only 16% of the organoids [24]. These models clustered tightly with NEPC patient tumors by transcriptomic and epigenomic profiling, expressed canonical NEPC signature genes including *MYCN*, *EZH2*, *SOX2*, *BRN2*, and *FOXA2*, and lacked AR expression and AR signaling activity [24].

Despite these advances, PDO models derived specifically from bone metastases of NEPC remain exceedingly rare. Most existing organoid lines originate from soft tissue or lymph node biopsies, and the biological and technical challenges of establishing organoids from bone metastasis have received limited attention. Bone represents the dominant site of lethal PC spread, and the bone microenvironment creates a distinct niche that shapes tumor cell behavior and therapeutic response [25–27]. Rapid autopsy programs (RAPs) provide a unique opportunity to harvest high-quality, freshly procured tumor tissue from multiple metastatic sites in patients who die from treatment-emergent disease. This offers an opportunity to study the biology of lethal CRPC, which is otherwise inaccessible through conventional biopsy. Here, we report the generation and comprehensive molecular characterization of patient-derived models of treatment-emergent NEPC established from bone and visceral metastases obtained through the PC RAP. By capturing the molecular features of end-stage metastatic disease, these models provide a clinically relevant and scalable platform for mechanistic studies and therapeutic discovery in treatment-emergent NEPC.

## RESULTS

### The Prostate Cancer RAP A Volunteer-Driven Tissue Procurement Program

To develop novel preclinical models of treatment resistant PC, we conducted a collaborative, multi-laboratory study with the PC RAP at the Fred and Pamela Buffett Cancer Center (FPBCC). The program was established in 2019 based on the foundational framework of the internationally renowned UNMC pancreatic cancer RAP [28]. The PC RAP is part of a broader tissue donation initiative at UNMC called the H.O.P.E. (Honoring Our Patients Every Day) program, which allows patients to contribute to cancer research posthumously through tissue donation. Supported by UNMC Institutional Review Board IRB#091-01-EP and IRB#460-19-EP, this infrastructure enables the systematic biobanking for tumor and unaffected tissues from hepatopancreatobiliary and PC donors. In collaboration with the regional organ procurement organization, Live On Nebraska, non-cancer tissues are also collected and archived by the H.O.P.E program. Biospecimens obtained through this resource have advanced research in early detection, imaging, therapeutic resistance, metastasis, genetics, and cachexia. **From Bedside to Bench:** The PC RAP workflow begins months to years before death, with enrollment of patients with advanced prostate cancer under the IRB 460-19-EP approval. When an enrolled donor enters palliative care, the GU oncology, pathology, and our laboratory teams are placed on alert to execute an autopsy within a critical 1-3 hour window after death. During the autopsy, specimens are barcoded and preserved using multiple complementary approaches, including snap-freezing for multi-omics, FFPE for histopathology, cryopreservation of viable tissue, and immediate initiation of patient-derived xenograft (PDX) and organoid (PDO) cultures.

### Establishment of Novel Preclinical Models from Metastatic CRPC

Figure 1A illustrates the disease trajectory of the donor patient from whom novel preclinical models were established. The deceased patient was diagnosed in 2019 with locally advanced prostate adenocarcinoma (Grade group 5) and metastasis to regional lymph nodes and bone. Shortly after, androgen deprivation therapy (ADT) was initiated with the LHRH agonist leuprolide in combination with abiraterone, a CYP17A1 inhibitor that blocks androgen biosynthesis, and the corticosteroid prednisone. Despite an initial response to therapy, the disease progressed and restaging revealed castration resistance and histopathological transformation to small cell neuroendocrine PC (SC-NEPC) with involvement of lymph nodes, bone, and visceral organs, including liver and lung. The patient subsequently received multiple lines of therapy, including carboplatin, docetaxel, cisplatin, etoposide, and radiotherapy, as well as the immune checkpoint inhibitor nivolumab. Despite these interventions, the patient succumbed to the disease in late 2021, approximately two years after diagnosis. This rapid disease progression highlights the aggressive, treatment resistant characteristics of lethal NEPC [29, 30].

**Figure 1.**
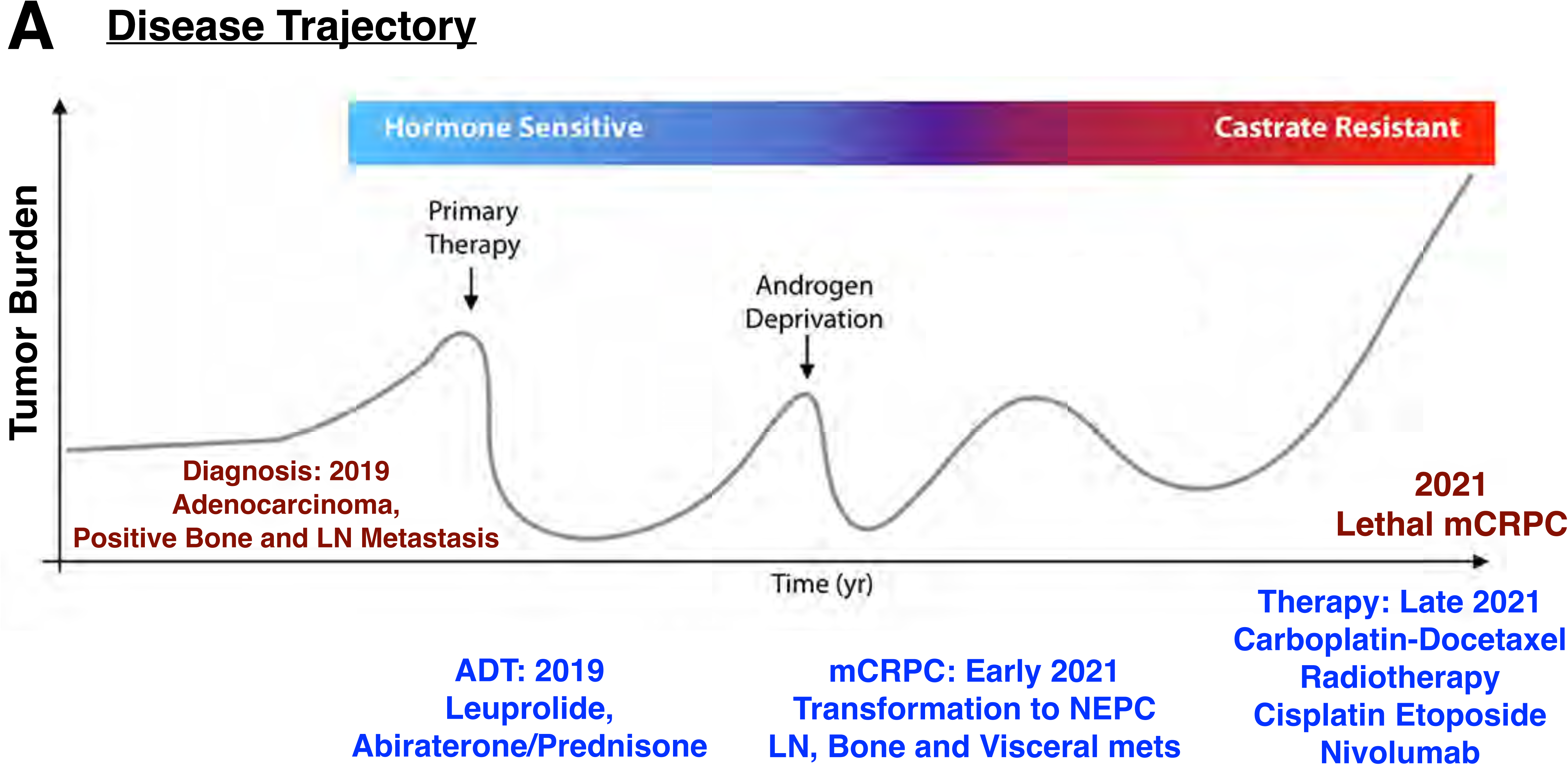
Disease trajectory of the PC RAP donor patient. (A) Schematic illustration of the clinical disease trajectory of the de-identified source patient (2021_UNMC_PRAD_RAP_2), from whom patient-derived preclinical models were established. The diagram depicts tumor burden over time, spanning the hormone sensitive and castration resistant timeline of the disease. The patient was diagnosed in 2019 with locally advanced prostate adenocarcinoma with regional lymph node and bone metastases and was initiated on ADT comprising Leuprolide acetate and Abiraterone/ Prednisone. He developed castration resistant transformation to small cell neuroendocrine PC with lymph node, bone, and visceral metastases in early 2021. Subsequent therapy included Carboplatin, Docetaxel, Cisplatin, Etoposide, Radiotherapy, and the anti PD-1 immunotherapy with Nivolumab. The patient succumbed to the disease in late 2021.

At autopsy, widespread metastatic involvement was observed, with disease in the liver, bone, lungs (right upper and lower lobes), lymph nodes, perinephric fat, and peripancreatic fat. Gross examination of the primary prostate revealed focal adenocarcinoma with therapy related changes, indicative of prior treatment effect. Liver metastases showed extensive parenchymal involvement (90%), with prominent gland formation (20%) and areas of necrosis (20%), consistent with a mixed neuroendocrine-adenocarcinoma phenotype. Bone marrow sections demonstrated severely dysplastic large cells infiltrating the marrow space, consistent with the osteolytic pattern characteristic of NEPC bone metastasis [31–33]. Histopathological analysis of procured tissue confirmed metastatic small cell carcinoma with neuroendocrine features (SC-NEPC) and an admixture of adenocarcinoma (CRPC-AD). This data is consistent with the well documented histological heterogeneity of lethal PC and the mixed lineage phenotype increasingly observed in patients with treatment-emergent CRPC NEPC [6, 29, 34]. Tissue obtained from the rapid autopsy was subsequently used to establish preclinical models following a standardized multi-modal workflow, as shown in Figure 2A. Each specimen was divided into five parallel processing streams: 1) fixation and paraffin embedding (FFPE) for histopathology; 2) snap-frozen for omics analyses; 3) cryopreservation of fresh viable chunks in 20% DMSO [35, 36]; 4) generation of patient-derived xenografts (PDX); and 5) collagenase digestion for generation of 2D cell lines, 3D organoids, and patient-derived organoid xenografts (PDOX).

**Figure 2.**
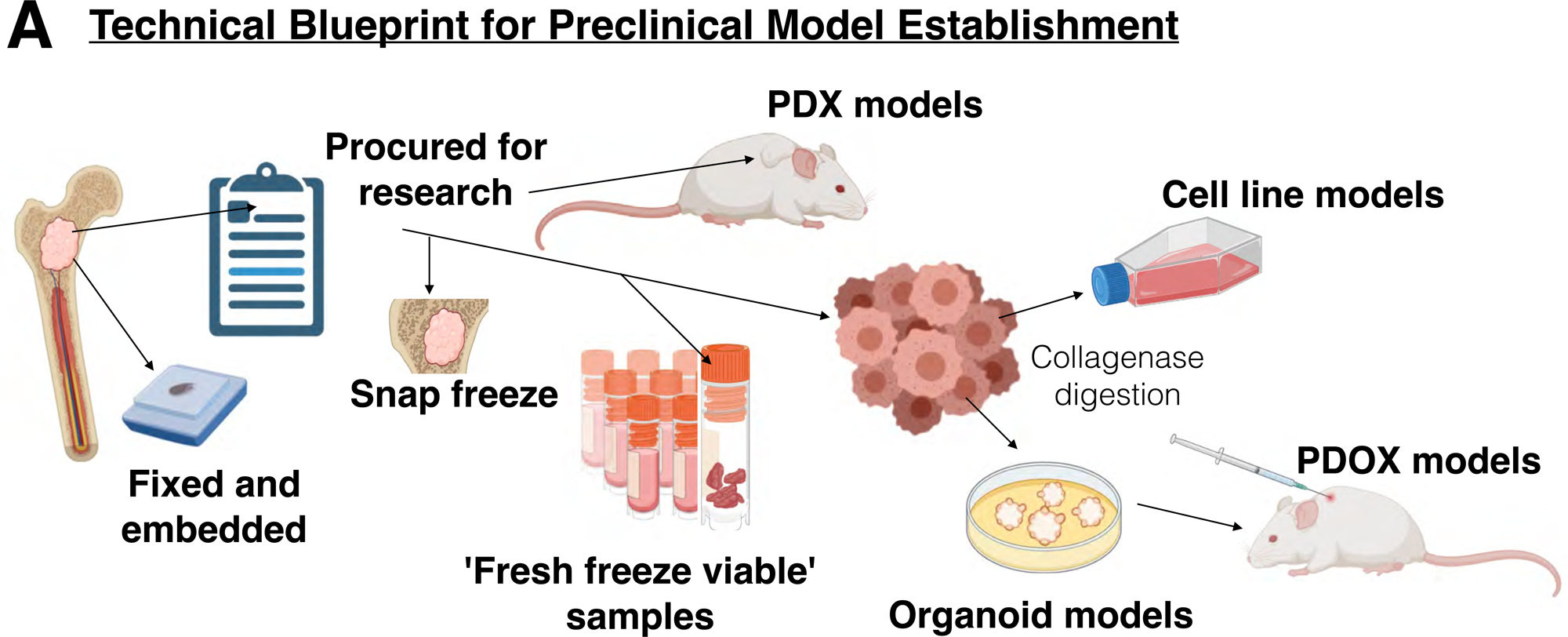
A technical blueprint for preclinical model establishment from PC RAP tissue. (A) Schematic overview of the standardized multi-step tissue processing workflow developed for preclinical model generation from PC RAP specimens. Upon receipt in the laboratory, each tissue sample was divided into five parallel processing streams: fixation and paraffin embedding (FFPE) for histopathological evaluation; snap freezing for multi-omics analyses; cryopreservation of fresh viable tissue chunks (1-2 mm) in 20% DMSO; collagenase digestion for the generation of 2D cell lines, 3D organoid models (PDO); and tissue chunks for establishing patient-derived xenografts (PDX).

### Genomic Profiling of PC RAP Tissue Identifies Treatment-Emergent NEPC Signatures

Through a long-standing collaborative relationship with the National Cancer Institute’s Patient-Derived Models Repository (NCI PDMR) focused on preclinical model development from rapid autopsies, viable specimens from the donor patient liver metastases were submitted to the NCI PDMR. Subsequently, through independent analysis of NCI PDMR datasets, two separate models were identified as matching UNMC PC RAP donor #2, and the corresponding banked and sequenced tissue specimens (331-R2_ORIGINATOR and 331-R4_ORIGINATOR) were analyzed. Publicly available whole exome sequencing (WES) data was used for comparative genomic analysis. OncoKB analysis of the PDMR WES data identified clinically relevant somatic alterations and a mutational landscape consistent with the genomic features of treatment-emergent NEPC, as summarized in Supplementary Table 1 [6, 29]. Shared mutations were detected in *ARID1A*, *TP53*, *ATM*, and *BRCA1* genes across both samples. A frameshift insertion in *ARID1A* (c.6267dup, p.H2090Tfs*9) was identified in both samples with variant allele frequencies (VAF) of 0.41-0.47, classified as likely loss-of-function. The tumor suppressor gene *TP53* harbored the same missense mutation (c.818G>A, p.R273H) in both samples with high VAF (0.95-0.99), consistent with a clonal driver mutation and classified as oncogenic with confirmed loss-of-function. Mutations in the DNA damage response gene *ATM* (c.998C>T, p.S333F) were present in both samples (VAF 0.45–0.47) but had inconclusive oncogenic and functional significance. *BRCA1* mutations (c.4535G>T, p.S1512I) were identified in both samples with lower VAF (0.30-0.33) and classified as likely neutral. Sample “331-R4_ORIGINATOR” harbored an additional *ZNF292* nonsense mutation (c.388C>T, p.Q130*) with high VAF (0.98), classified as likely oncogenic and loss-of-function. Variant analysis revealed 6 missense mutations (67%), 2 frameshift insertions (22%), and 1 nonsense mutation (11%). Mean VAF across all variants was 0.595 (median 0.466, range 0.30-0.99), with clonal mutations (*TP53, ZNF292*) displaying VAF >0.90 and subclonal variants (*BRCA1, ATM*) showing VAF <0.50.

To further characterize the genomic landscape of the source tumor, we re-analyzed the PDMR WES FASTQ files to assess ploidy and genome-wide copy number alterations, see Figure 3A-C. Analysis of the two originator tumor samples (331-R2 and 331-R4) revealed a complex copy number landscape with widespread chromosomal alterations across the genome, including regions of amplification, copy loss, and loss of heterozygosity (LOH). Notably, we observed copy-neutral LOH on chromosome 10 encompassing the *PTEN* locus and on chromosome 13 encompassing the *RB1 and BRCA2* loci in both originator samples (Figure 3A-B). The B-allele frequency plot further confirmed allelic imbalance at these loci, consistent with LOH rather than homozygous deletion (Figure 3B).

**Figure 3.**
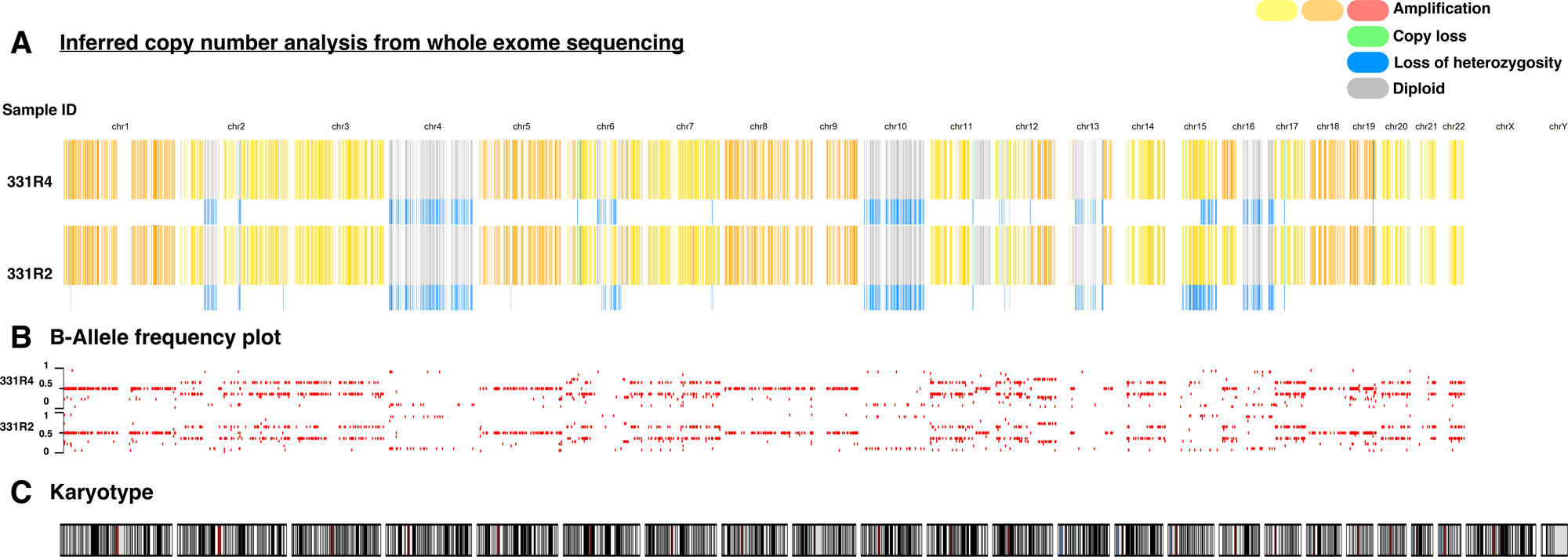
Genomic characterization of PC RAP source tumor reveals copy number alterations consistent with treatment-emergent NEPC. (A) Inferred copy number analysis from whole exome sequencing (WES) of two originator tumor samples (331-R4 and 331-R2) across all autosomes and sex chromosomes. Copy number states are color-coded: amplification (red), copy loss (green), loss of heterozygosity (blue), diploid (grey), and gains (orange/yellow). Notable regions of loss of heterozygosity are at chromosome 10 (*PTEN* locus) and chromosome 13 (*RB1* locus). (B) B-allele frequency (BAF) plot for samples 331-R4 and 331-R2 across all chromosomes, confirming allelic imbalance consistent with loss of heterozygosity at key tumor suppressor loci. BAF values range from 0 to 1. (C) Karyotype derived from copy number analysis. WES data were obtained from the NCI Patient-Derived Models Repository (PDMR) and re-analyzed for ploidy and copy number alteration assessment.

### Bone and Liver Metastasis Derived Cellular Models from Rapid Autopsy Retain Molecular Features of Lethal NEPC

A central goal of our PC RAP based modeling effort was to derive cellular models that preserve the histopathological and molecular features of the metastatic tissue from which they were established. To assess this, we systematically compared the morphological and immunophenotypic characteristics of the cell models with matched patient tissue procured at autopsy. We interpreted our findings in the context of the established clinicopathological literature on bone and liver metastases in lethal NEPC. Although WES was not performed on our PC RAP derived cellular models, the genomic data presented above reflect the molecular profile of the source patient tissue and provide a close representation.

SC-NEPC was first described in the context of ectopic adrenocorticotropic hormone production and paraneoplastic syndrome [37]. The clinicopathological features of SC-NEPC include uniform expression of synaptophysin, chromogranin, and high/low molecular weight cytokeratins, with high Ki-67 labeling indices (≥70%) [31, 32, 38, 39]. Most tumors lose the prostate specific transcription factor NKX3.1, and about half gain the lung/thyroid specific transcription factor TTF-1, a pattern typically associated with small-cell lung cancer [39]. Liver metastasis in NEPC is a hallmark of advanced, treatment-emergent disease and an independent poor prognostic indicator alongside *RB1/ TP53* loss and low PSA [30]. Histopathological assessment of the liver metastases from the RAP patient demonstrated a high grade tumor with mixed features of both SC-NEPC and focal gland formation as seen in adenocarcinoma (Figure 4A, H&E). Next, we performed immunofluorescence (IF) staining of the liver metastases across a panel of six markers: cytokeratin 8/18, KRT8/18; neural cell adhesion molecule, NCAM1; marker of proliferation, MKI-67; synaptophysin, SYP; Enhancer of Zeste Homolog 2, EZH2; and NK3 homeobox 1, NKX3.1. Consistent with published literature, IF staining revealed strong expression of Cytokeratin 8/18, confirming prostate epithelial lineage, alongside robust NCAM1 expression and a high Ki-67 labeling index, reflecting the aggressive proliferative nature of the mixed neuroendocrine-adenocarcinoma tumors (Figure 4A and whole tissue scan in Supplemental Figure 1). We observed moderate positivity within the tumor for Synaptophysin (SYP), with signal concentrated in discrete cell clusters (Supplemental Figures 1&2). This focal SYP expression pattern is consistent with the variable, often patchy, chromogranin A and synaptophysin staining observed in poorly differentiated neuroendocrine carcinomas and mixed histology tumors [40, 41]. We observed expression of EZH2 throughout the tissue (Supplemental Figures 1&2), confirming that EZH2 upregulation is a likely cancer cell-intrinsic feature of the liver metastasis tissue. This observation is consistent with EZH2 overexpression reported across CRPC-NE patient tumors [24, 42, 43]. On the contrary, NKX3.1 was markedly absent throughout the tissue, with no appreciable nuclear immunoreactivity in tumor cells; see Supplemental Figures 1&2. This confirms the loss of AR regulated luminal transcription factor activity at the cellular level and is consistent with the downregulation of AR expression in treatment-emergent NEPC [6].

**Figure 4.**
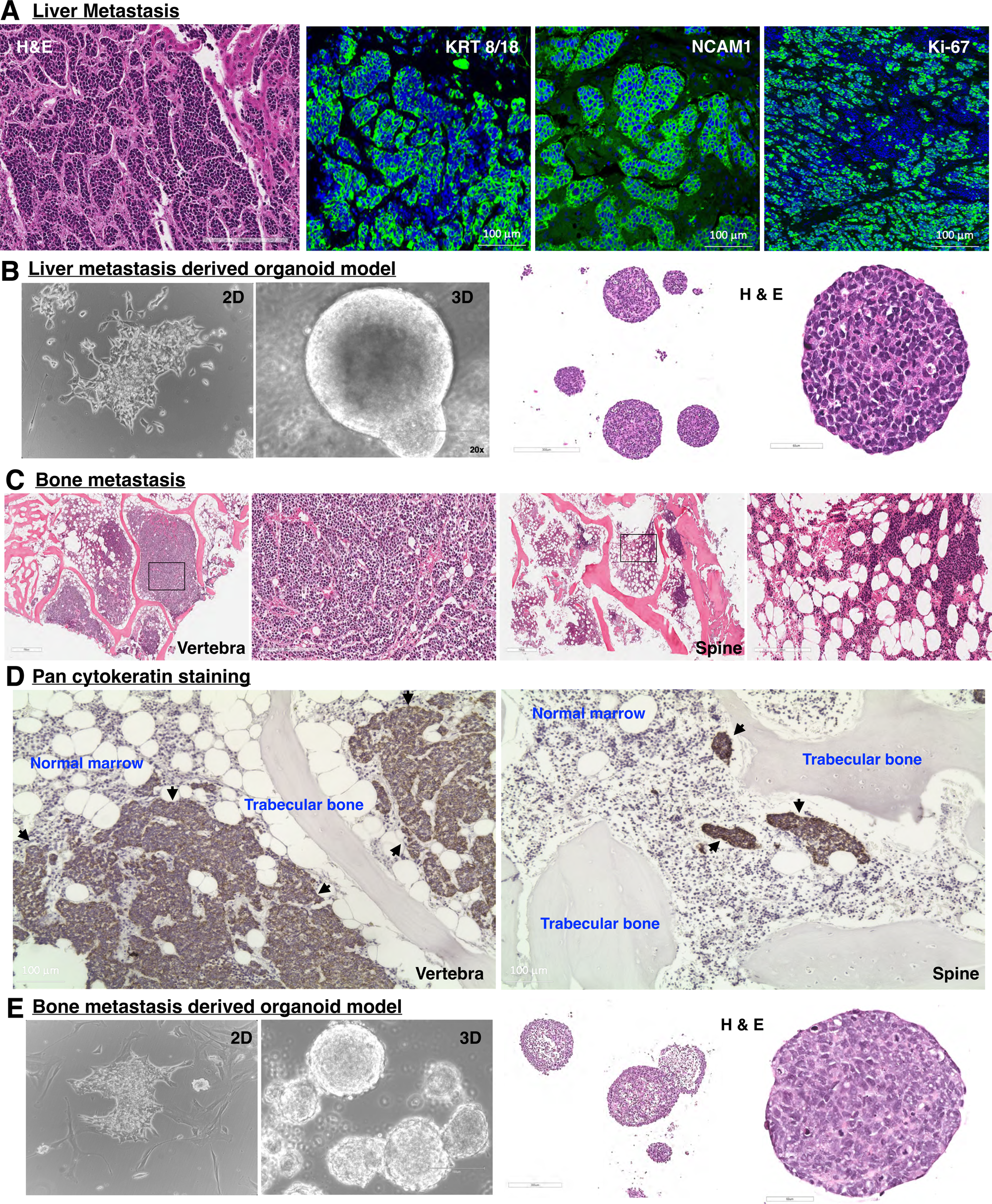
Histopathological characterization of PC RAP liver, bone metastases, and derived organoid models. (A) Histopathological and immunofluorescence characterization of liver metastasis tissue, showing H&E staining (left; scale bar 200 µm) and multiplex immunofluorescence for Cytokeratin 8/18 (CK8/18), NCAM1, and Ki-67 (green), with DAPI nuclear counterstain (blue), scale bar 100 µm. (B) Representative bright-field images of PDO_LMO in 2D monolayer (left) and 3D organoid (middle; scale bar, 150 µm) culture at passage ≥10, with corresponding H&E stained organoid sections at low (left; scale bar, 300 µm) and high (right; scale bar, 60 µm) magnification. Scale bars, 300 µm. (C) Hematoxylin and eosin (H&E) staining of bone metastasis tissue procured from vertebral (left) and spinal (right) sites at low magnification (left panels; scale bar 700 µm) with corresponding high magnification insets (right panels; scale bar 200 µm) demonstrating tumor infiltration of the bone marrow space with small to medium sized cancer. (D) Pan-cytokeratin immunohistochemical staining of vertebral (left) and spinal (right) bone metastasis sections, confirming the epithelial identity of infiltrating tumor cells (arrows) within normal marrow and trabecular bone architecture. Scale bars, 100 µm. (E) Representative bright-field images of PDO_BMO in 2D monolayer (left) and 3D organoid (middle; scale bar 150 µm) culture at passage ≥10, with corresponding H&E-stained organoid sections at low (left; scale bar, 300 µm) and high (right; scale bar, 60 µm) magnification demonstrating solid spheroid architecture with high nuclear-to-cytoplasmic ratios and frequent mitotic figures.

Next, we performed dual IF staining with a panel of epithelial differentiation (KRT8/18, CDH1), neuroendocrine differentiation (SYP, NCAM1), cell proliferation (Ki-67), and mesenchymal plasticity (vimentin, VIM) markers. We observed co-expression of KRT8/18 and NCAM1 and EZH2, with strong NCAM1/EZH2 staining in KRT8/18 positive cells, indicating active neuroendocrine differentiation and confirming the dual epithelial-neuroendocrine phenotype characteristic of treatment-emergent CRPC-NE (Supplementary Figure 3). Ki-67 staining revealed a proliferative population within these NEPC cells, with retained CDH1 expression suggesting maintenance of epithelial adhesion despite neuroendocrine transdifferentiation (Supplementary Figure 3). The expression of vimentin was mostly localized to blood vessels within the tissue (Supplementary Figure 3). Our data confirm that the originating autopsy tissue exhibits the characteristic molecular hallmarks of CRPC-NE with active proliferation.

Unlike the predominantly osteoblastic lesions of prostate adenocarcinoma, bone metastases in NEPC are characteristically osteolytic [Sharpe & McDonald, Arch Pathol 1942 & 44, 45-47]. The histopathological evaluation of bone metastasis tissue procured from vertebral and spinal sites of the source patient revealed extensive tumor infiltration of the bone marrow space, with diffuse sheets of small to medium sized malignant cells. These cells (Figure 4C) displayed nuclear molding, granular chromatin pattern, inconspicuous nucleoli, and frequent mitotic figures, all of which are diagnostic of SC-NEPC [40, 48–51]. Pan-cytokeratin immunostaining confirmed the epithelial identity of the infiltrating tumor cells, which were clearly distinguishable from surrounding normal marrow and trabecular bone at both vertebral and spinal sites. We observed cytokeratin-positive tumor cell clusters extending diffusely within the trabecular bone (Figure 4D, arrows).

### Establishing Translationally Relevant Cellular Models of NEPC

Prior work has demonstrated successful long-term culture of PC organoids from patient biopsies [21], a 16% success rate (4/25) for NEPC PDO generation from fresh metastatic tissue [24], and long-term maintenance of over 50% of CRPC PDX-derived organoid lines with conserved genomic and lineage features across passages [13]. We obtained fresh tissue chunks from the primary prostate tumor and three metastatic sites (lung, liver, and bone) to establish both 2D cell lines and 3D organoid models from each sample. We followed previously published organoid generation protocols [52–56]. The prostate derived cells showed initial growth but did not survive beyond passage 2, while lung-derived tissue did not generate viable cultures. In contrast, cells derived from liver and bone metastases grew consistently as both 2D monolayer and 3D organoids and were successfully passaged beyond passage 10 (Figure 4B&E, phase-contrast images). In 2D culture, the liver and bone metastasis-derived cells displayed a small-cell morphology consistent with the neuroendocrine phenotype of the source tumor. In 3D culture, both liver and bone metastases derived organoid lines, LMO and BMO, respectively, formed well-defined spheroids of variable size. H&E staining of organoid sections (Figure 4B&E, H&E) revealed solid spheroid architecture composed of tightly packed cells with high nuclear-to-cytoplasmic ratios, nuclear molding, and frequent mitotic figures, closely recapitulating the small cell morphology observed in the source patient tissue (Figure 4A&C, H&E). The morphological concordance between patient tumor and derived organoids aligns with prior NEPC studies and confirms model fidelity to the tissue of origin [6, 7]. Overall, we achieved a 50% success rate for long-term preclinical model development from PC RAP procured tissue. This is comparable to the PDX to organoid culture rates reported in the LuCaP cohort [13] and substantially higher than the 10 to 16% success rates typically achieved from CRPC needle biopsies [21]. Notably, patient-derived organoids (PDOs) that sustain growth past passage 10 are generally maintainable long-term in culture, as supported by published data [13].

### PC Bone Metastasis-Derived Mesenchymal Stem Cells (MSCs) Express Classical MSC Markers but Display Reduced Osteogenic Potential

Advanced PC tumors commonly metastasize to bone and induce pathologic bone formation, resulting in weak, brittle bone that is prone to fracture and, as a result, poor patient quality of life. PC-induced bone disease has been identified to be driven, in part, by cancer cell interactions and regulation of MSCs in the bone environment [57–59]. Several studies have examined primary and metastasis-derived PC cells; however, we wanted to examine the phenotype of MSCs from the tumor-bone environment. To do this, the patient’s spinal fragments, adjacent to metastases, were dissociated *ex vivo* and the cell supernatant was expanded in standard MSC culture conditions. The resultant cell populations exhibited mesenchymal/fibroblastic cell morphology (Figure 5A, left). Given their morphology and tissue source, we hypothesized that the cells were MSCs. We therefore performed flow cytometry to examine cell surface MSC marker expression. The RAP-derived cells displayed high expression of MSC markers CD90, CD105, and CD73 (Figure 5A, right) [60]. In contrast, there was an absence of hematopoietic cell markers CD45, CD34, CD11b, CD19, and HLA-DR [60].

**Figure 5.**
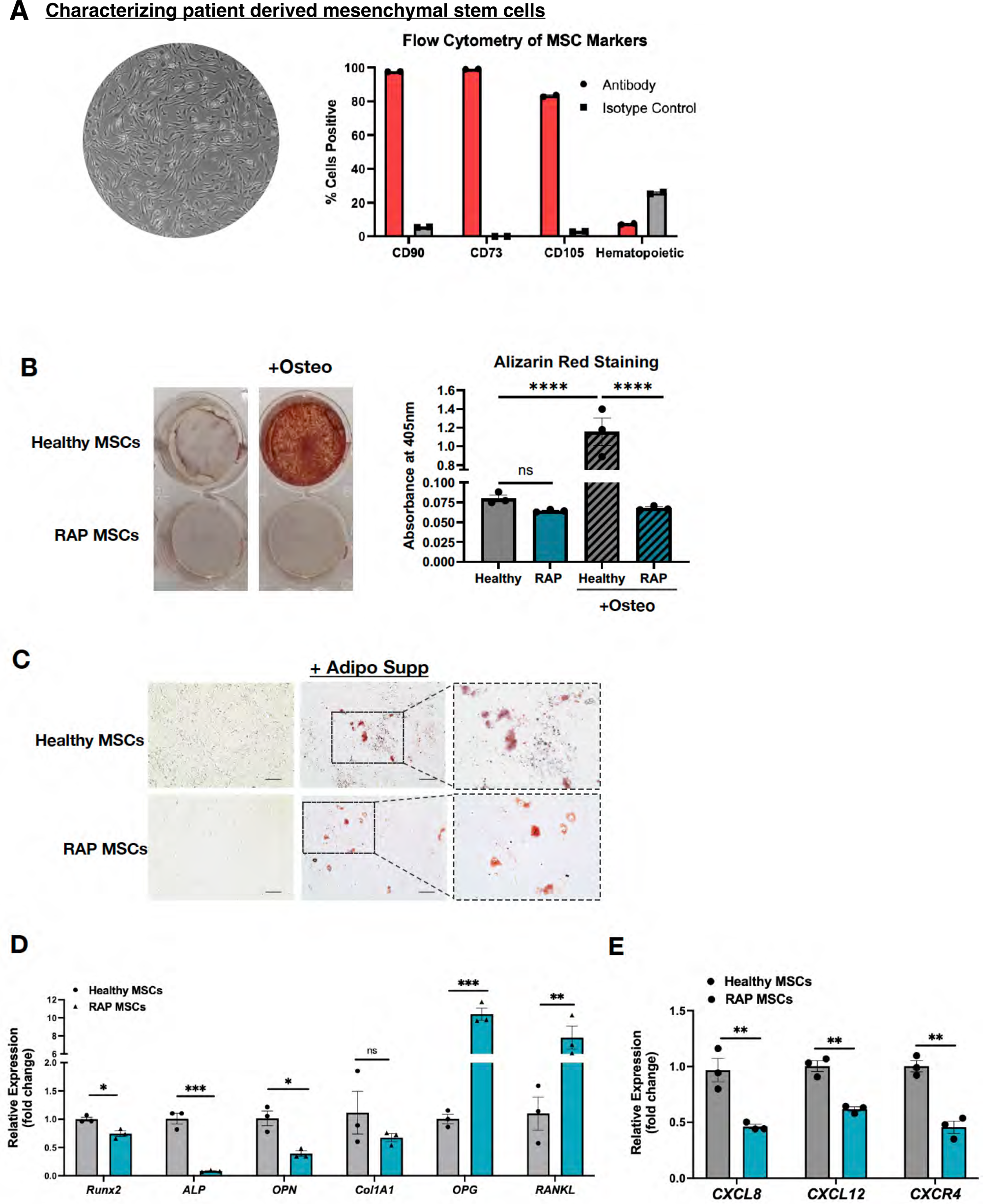
Characterization of bone metastasis-derived mesenchymal stem cells (MSCs). (A) Phase contrast image of RAP MSCs (left); flow cytometry of MSC markers (CD90, CD105, CD73) and hematopoietic cell markers (CD45, CD34, CD11b, CD19, HLA-DR) (right); graph shows percentage of viable cells; (B) Osteogenesis differentiation assay; representative images of calcium deposition (Alizarin Red stain) in healthy donor and RAP MSCs following treatment of osteogenic differentiation supplement (+Osteo) (left). Calcium deposits were quantified using alizarin red staining; the representative graph shows absorbance of alizarin red after acetic acid extraction from each well (right). (C) Adipogenesis assay; representative brightfield microscope images of Oil Red O stained healthy donor and RAP MSCs following treatment with adipogenic differentiation supplement (+Adipo). Scale bar, 100 µm; (D) Real-Time qPCR of RAP MSCs compared with 2 different sources of bone-derived “healthy” MSCs. The graph shows the expression of genes associated with osteogenesis. (E) Real-time qPCR of RAP and healthy MSCs; the graph shows gene expression of chemokines and chemokine receptors. Graphs represent relative fold change of genes normalized to 36B4 rRNA expression. *p < 0.05, **p < 0.01, ***p < 0.001, and ****p < 0.0001 one-way ANOVA followed by Dunnett’s multiple comparison test.

There has been substantial evidence to demonstrate that metastatic PC promotes osteoblast differentiation from bone-resident MSCs [61–64]. In order to understand the impact of advanced PC on the phenotype of MSCs isolated from the RAP sample, we performed differentiation assays for comparison to bone MSCs from healthy donors. To examine the potential for osteogenic differentiation, MSCs were treated with osteogenesis-inducing supplement. Using Alizarin Red staining as a read-out for matrix mineralization, there was visibly more calcium deposition by healthy donor MSCs compared to little to no visible calcium deposition by RAP MSCs (Figure 5B, left). Quantitation of these data revealed that the osteoblastic differentiation of RAP MSCs was significantly impaired compared to MSCs from normal bone, which exhibited ∼16-fold increase in calcium deposition compared to RAP MSCs (Figure 5B, right). Given the complete loss of the RAP MSC osteoblastic differentiation ability, we considered the possibility that these RAP MSCs had lost all differentiation capacity. To test this idea, we treated healthy donor and RAP MSCs with adipogenesis-inducing supplement and measured lipid droplets using Oil Red O stain. In contrast to osteogenesis potential, there was no difference in adipogenesis of RAP MSCs, compared to healthy donor MSCs (Figure 5C). Based on an observable inability to differentiate into osteoid-generating cells, we next examined expression of genes associated with osteogenesis in RAP MSCs. Runx2 is the major osteoblast differentiation transcription factor that activates expression of other osteogenesis genes [65]; similar to the differentiation assay, *Runx2* expression was significantly less in RAP MSCs compared to healthy donor MSCs (Figure 5D). We saw similar and highly significant reduction in Runx2 downstream target genes, alkaline phosphatase (ALP), osteopontin (OPN), and type I collagen (Col1a1), which each contribute to bone matrix mineralization albeit in a different capacity [66, 67]. In coupled bone remodeling (i.e., balanced osteogenesis/bone formation and osteolysis/ bone resorption), MSCs and precursor osteoblasts produce receptor activator of nuclear factor Kappa-Β ligand (RANKL), which induces osteoclast formation, and its inhibitor, decoy receptor osteoprotegerin (OPG) [68]. Decreased expression of osteogenesis-associated genes suggests a decreased ratio of osteogenesis to osteolysis. Thus, we examined MSC expression of *RANKL* and *OPG*. Interestingly, there was a significant increase in RANKL expression (∼8-fold) in RAP MSCs compared to healthy donor MSCs. However, there was also a significant increase in RAP MSC *OPG* expression (Figure 5D). Last, based on evidence demonstrating important roles for chemokine signaling in MSCs and the tumor-bone microenvironment [69, 70]. We measured expression of specific chemokines, *CXCL8*, *CXCL12*, and *CXCR4* (CXCL12 receptor) in RAP-derived MSCs compared to healthy MSCs. We observed that RAP MSCs express significantly less CXCL8 and CXCL12, as well as CXCR4 suggesting that there would be less chemokine-mediated contribution to the tumor in bone compared to healthy MSCs (Figure 5E). However, these chemokines also contribute to immune regulation and further experiments would be necessary to delineate the importance of these changes in the tumor-bone interface. Overall, gene expression analysis in the RAP MSCs shows a decrease in many essential osteogenesis genes and provides further evidence of an impairment of these cells’ ability to differentiate into osteoblasts.

### Transcriptomic Profiling of RAP-Derived Organoid Models Matches the Molecular Identity of NEPC

To assess whether PDO lines recapitulate the source tumor transcriptome, we performed bulk RNA sequencing on bone (PDO_BMO) and liver (PDO_LMO) metastasis-derived organoids and compared their expression profiles to normal prostate, primary PC, and metastatic PC (mPC) samples from the NCI Genomic Data Commons portal. Principal component analysis (PCA) showed that both PDOs (LMO1/LMO2 and BMO1/BMO2 biological replicates, passage 10) clustered tightly within the mPC group and separated from normal prostate and primary tumor samples along PC1 (38% of total variance; Figure 6A). Notably, the PDO lines clustered closely with NCI-H660, the only established and commercially available human NEPC cell line, and were distinct from AR driven adenocarcinoma lines such as LNCaP (Figure 6B). This pattern is consistent with their neuroendocrine molecular identity and mirrors the transcriptomic clustering patterns previously reported for patient-derived NEPC organoids [24]. Unsupervised hierarchical clustering of differentially expressed genes further confirmed this NEPC identity. The NEPC cluster was characterized by high expression of canonical neuroendocrine and lineage plasticity genes (Figure 6B, asterisk), including *AURKA*, *MYCN*, *TP53*, *NCAM1*, *SOX2*, *PROX1*, and *SYT11*, and low expression of *AR* driven luminal genes, including *AR*, *KLK3*, *KLK4*, and *NKX3-1*, consistent with the transcriptomic signature of treatment emergent NEPC described in landmark molecular profiling studies [6, 71–73]. Downregulation of *RB1* and *PTEN* was also evident in the PDO lines, in keeping with the frequent co-occurrence of *RB1* and *TP53* loss as drivers of lineage plasticity in NEPC [24, 74, 75].

**Figure 6.**
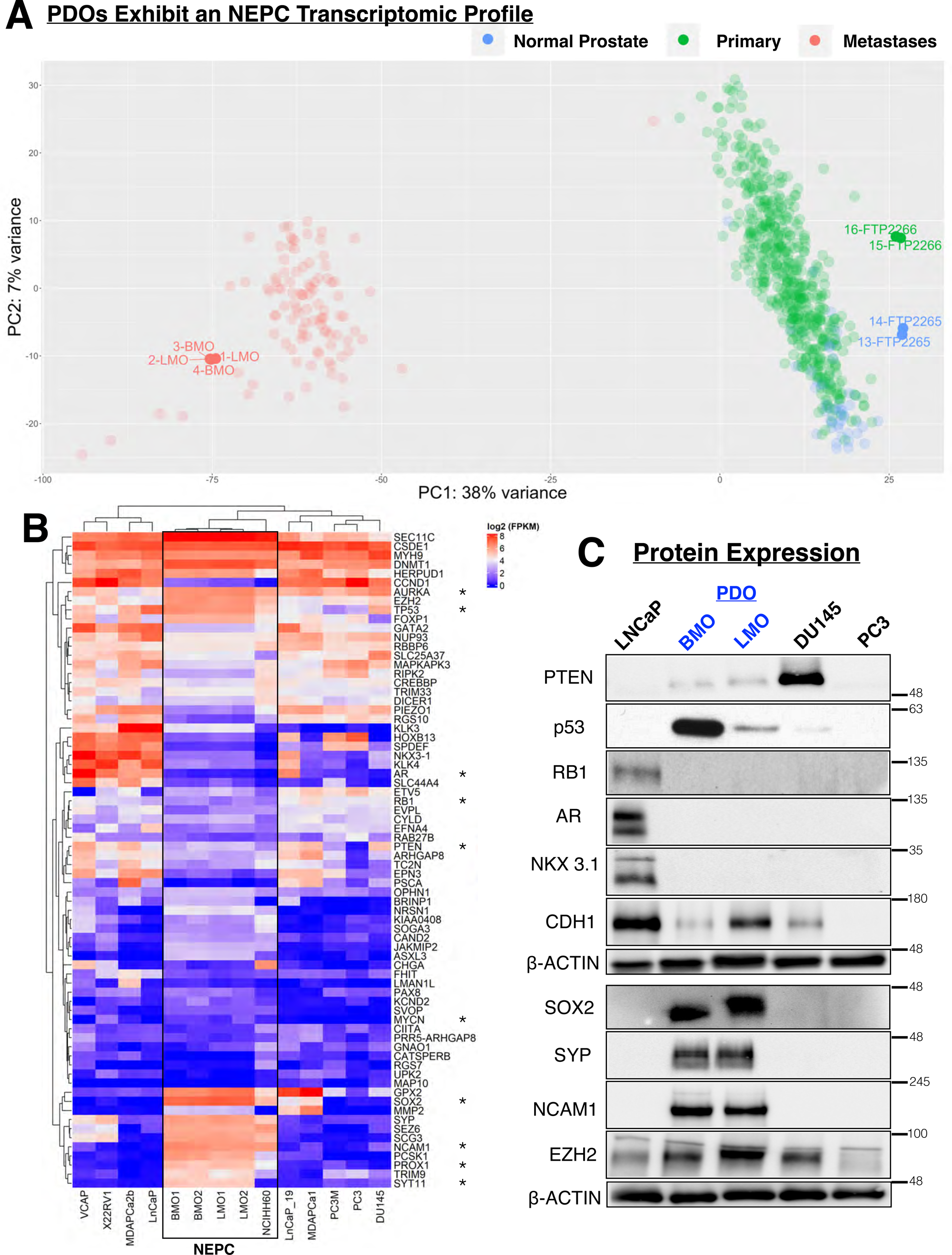
Transcriptomic profiling confirms NEPC molecular identity of PC RAP-derived organoid lines. (A) PCA of RNA sequencing data comparing PDO_BMO (passage 5 and passage 10) and PDO_LMO (passage 5 and passage 10) against a reference cohort of samples from data on the Genomic Data Commons (GDC) portal, comprising normal prostate (blue), primary PC (green), and metastatic PC (red). PC1 accounts for 38% of total variance; PC2 accounts for 7% of total variance. (B) Unsupervised hierarchical clustering heatmap of differentially expressed genes across PDO_BMO, PDO_LMO, and a panel of established PC cell lines. Gene expression values shown as log2 FPKM. The NEPC cluster is highlighted by a black box. (C) Western blot analysis of adenocarcinoma lineage markers (PTEN, TP53, RB1, AR, NKX3.1, CDH1) and neuroendocrine markers (SOX2, SYP, NCAM1, EZH2) in PDO_BMO and PDO_LMO compared to established PC cell lines (LNCaP, DU145, PC3). Beta-actin served as a loading control. Molecular weights are indicated in kilodaltons (kDa). AR, androgen receptor; CDH1, E-cadherin; EZH2, enhancer of zeste homolog 2; NCAM1, neural cell adhesion molecule 1; NKX3.1, NK3 homeobox 1; PTEN, phosphatase and tensin homolog; SOX2, SRY-box transcription factor 2; SYP, synaptophysin; TP53, tumor protein p53; RB1, retinoblastoma.

To validate gene expression findings at the protein level, we performed western blot analysis on the PDO lines alongside established metastatic PC cell lines LNCaP, DU145, and PC3 as reference models (Figure 6C). Consistent with their NEPC transcriptomic profile, both PDO lines showed markedly reduced or absent expression of the adenocarcinoma lineage markers AR, NKX3.1, and PTEN, and reduced expression of the epithelial adhesion molecule CDH1 relative to LNCaP. In contrast, both lines demonstrated robust expression of neuroendocrine markers SOX2, SYP, EZH2, and NCAM1. Interestingly, p53 protein levels differed between the two PDO lines. We observed robust p53 protein expression in the bone metastasis-derived PDO (PDO_BMO), whereas the liver metastasis-derived PDO (PDO_LMO) exhibited low p53 expression. This data potentially reflects intratumoral heterogeneity in *TP53* mutation type between the bone and liver metastatic sites. We observed reduced RB1 expression in both PDOs relative to the adenocarcinoma control LNCaP [29]. Importantly, our findings were concordant with the genomics data obtained from the PDMR originator tissue (Figure 3A and Supplemental Table 1). RB1 protein loss in both lines was consistent with loss of heterozygosity at the *RB1* locus on chromosome 13 identified by WES, while PTEN protein loss was concordant with copy-neutral loss of heterozygosity at chromosome 10 encompassing the *PTEN* locus [76, 77]. Taken together, the genomic, transcriptomic, and protein expression data confirm that PDO_LMO and PDO_BMO faithfully recapitulate the molecular identity of treatment-emergent NEPC and maintain it stably across multiple passages in culture [6, 24, 73, 74, 77, 78].

To determine whether the PDOs faithfully recapitulate the molecular phenotype of the source patient tumor, we performed comprehensive IF profiling of organoid sections across a panel of epithelial, neuroendocrine, stem cell, epigenetic, and proliferative markers (Figure 7). Both PDO lines retained strong expression of the luminal epithelial marker Cytokeratin 8/18 (KRT8/18), supporting preservation of prostate epithelial lineage identity across passages and consistent with prior reports of KRT8/18 retention in prostatic small cell carcinoma [47, 79]. Cytokeratin 5/14 (KRT 5/14) was detected in both lines. However, the staining pattern was diffuse and faint with cytoplasmic distribution rather than the discrete cell membrane localization characteristic of true basal epithelial cells, suggesting incomplete and aberrant retention of basal identity likely reflective of the plasticity permissive state of these tumors [75]. NKX3.1, a canonical AR regulated marker of luminal prostate epithelial differentiation, was similarly absent in both organoid lines, with no detectable nuclear staining, consistent with the loss of AR driven luminal identity in treatment emergent NEPC [6]. In contrast, both organoid lines expressed canonical neuroendocrine markers, supporting their CRPC-NE molecular identity. Synaptophysin (SYP) and NCAM1 were detectable in both lines, with particularly robust NCAM1 expression, while we observed moderate staining for Chromogranin A (CHGA), consistent with the variable and often focal CHGA expression documented in poorly differentiated neuroendocrine carcinomas [80, 81]. Strong nuclear expression of SOX2 was observed in both PDOs, consistent with its established role as a lineage plasticity driver and neural transcription factor in treatment emergent NEPC [75, 82–86]. Notably, both organoid lines demonstrated strong nuclear EZH2 expression, supporting a role for this epigenetic modifier in maintaining the neuroendocrine state and suppressing luminal differentiation programs in these models, consistent with EZH2 overexpression reported in the majority of CRPC-NE patient tumors and organoids [6, 24, 74]. Ki-67 staining confirmed active proliferation in both lines, with positive nuclei distributed throughout the organoid sections, reflecting the high proliferative index characteristic of SC-NEPC [30]. Taken together, the co-expression of luminal epithelial markers (KRT 8/18, KRT 5/14) alongside neuroendocrine markers (SYP, CHGA, NCAM1), neural transcription factors (SOX2), and epigenetic regulators (EZH2) in both bone and liver metastases derived PDOs (Figure 7) recapitulates the heterogeneous, plasticity permissive molecular phenotype of the PC-RAP patient tumor. This intermediate lineage pattern, characterized by retention of epithelial identity alongside acquisition of neuroendocrine features (CRPC-NE and AD admixture), is increasingly recognized as a defining feature of treatment-emergent CRPC-NE [75, 87].

**Figure 7.**
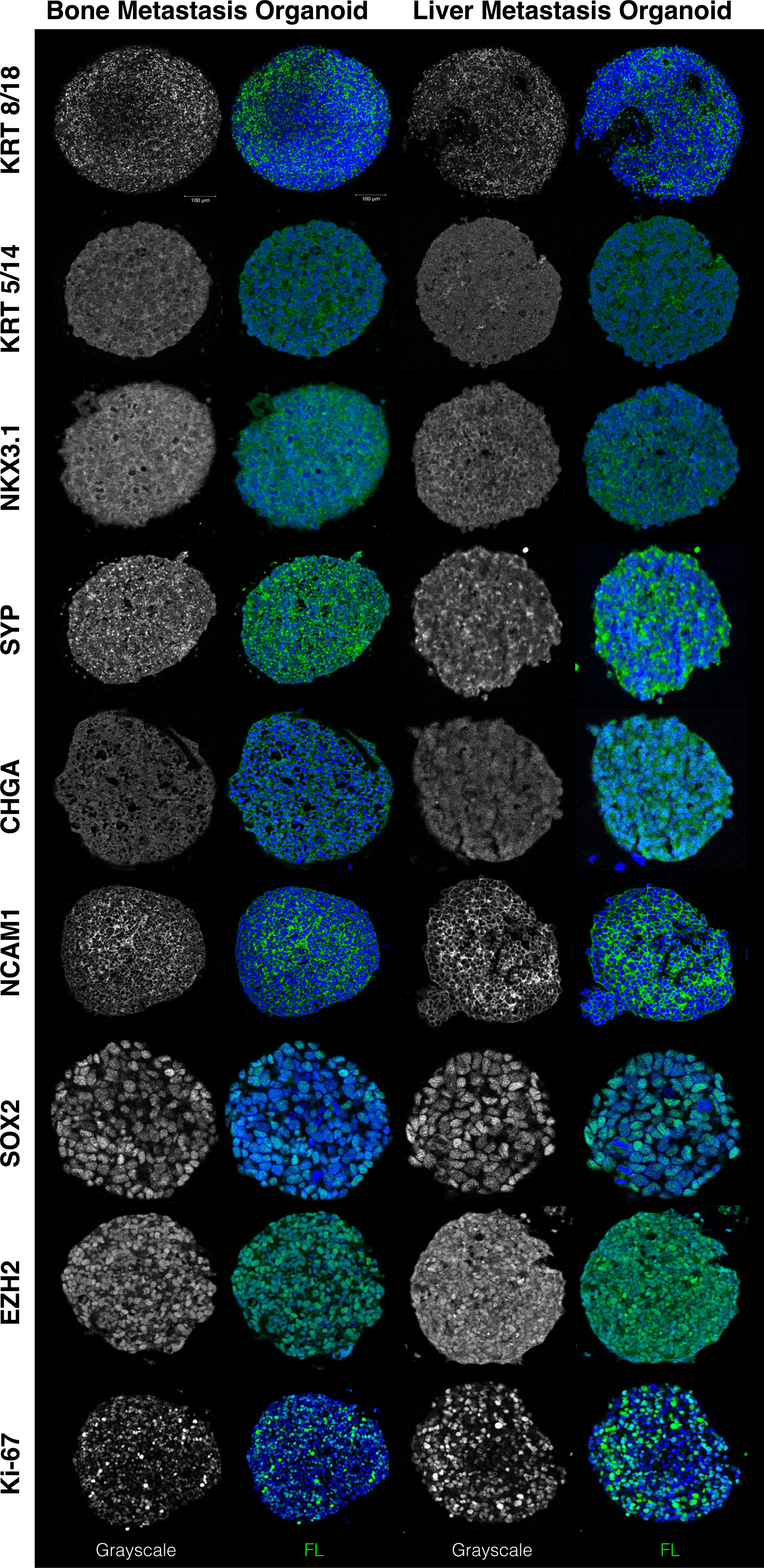
Comprehensive immunofluorescence profiling of PC RAP-derived organoids confirms plasticity permissive NEPC phenotype. Immunofluorescence staining of PDO_BMO (left two columns) and PDO_LMO (right two columns) organoid sections at passage ≥10 for a panel of epithelial, neuroendocrine, stem cell, epigenetic, and proliferative markers: Cytokeratin 8/18 (KRT 8/18), Cytokeratin 5/14 (KRT 5/14), NKX3.1, Synaptophysin (SYP), Chromogranin A (CHGA), NCAM1, SOX2, EZH2, and Ki-67. For each marker, grayscale (left) and pseudocolor merged (right) images are shown, with marker signal in green and DAPI nuclear counterstain in blue. Scale bars, 100 µm.

### PC RAP-Derived Cell Lines Exhibit an Elevated Antioxidant Defense Program

Radiation is a frequent treatment modality in mCRPC, and we therefore evaluated the radiosensitivity of the bone and liver metastasis derived cell lines as a proof-of-concept functional study. The radiation history of the harvested tissue was incompletely defined. Although the patient had received external beam radiotherapy to several bone metastatic sites, we could not determine whether the specific tissue samples harvested lay within a treated field. In contrast, the liver metastases were not irradiated in situ, and liver derived tissue can be considered radiation-naïve. Radiation exposure and hypoxic stress represent key features of the metastatic tumor microenvironment that can influence oxidative stress responses and treatment resistance [88–91]. Therefore, to assess how mCRPC patient derived cells respond to oxidative and microenvironmental stress, bone and liver metastasis derived cell lines were subjected to ionizing radiation (2 Gy in normoxia) or maintained under hypoxic conditions. Intracellular reactive oxygen species (ROS) generation was quantified using the ROS sensor/ fluorescent probes 2′,7′-dichlorodihydrofluorescein diacetate (DCF) and dihydroethidium (DHE). DCF was used as a general indicator of total reactive oxygen species (ROS) levels, detecting a broad range of oxidant species, whereas DHE provided a more specific measure of superoxide generation. The bone and liver metastasis-derived (Figure 8A-B,D-E and Supplementary Figure 4) cell lines showed significantly elevated total ROS and superoxide levels under 2 Gy radiation or hypoxia compared to untreated control. To investigate the molecular basis of this tolerance, we examined the expression of key antioxidant defense enzymes. Under hypoxic conditions, both cell lines (Figure 8C and F) maintained expression of the mitochondrial antioxidant MnSOD, and the cytoplasmic antioxidant CuZnSOD. The hydrogen peroxide scavenging enzyme, Catalase, increased in both cell lines and did so in response to radition or hypoxia. These findings are consistent with the well documented upregulation of antioxidant defense mechanisms in radioresistant cancer cells [92, 93]. Collectively, these data provide a proof-of-concept demonstration that the PC RAP derived cell lines can be used to model clinically relevant treatment resistance phenotypes, and the antioxidant defense pathway is a potential therapeutic vulnerability in mCRPC.

**Figure 8.**
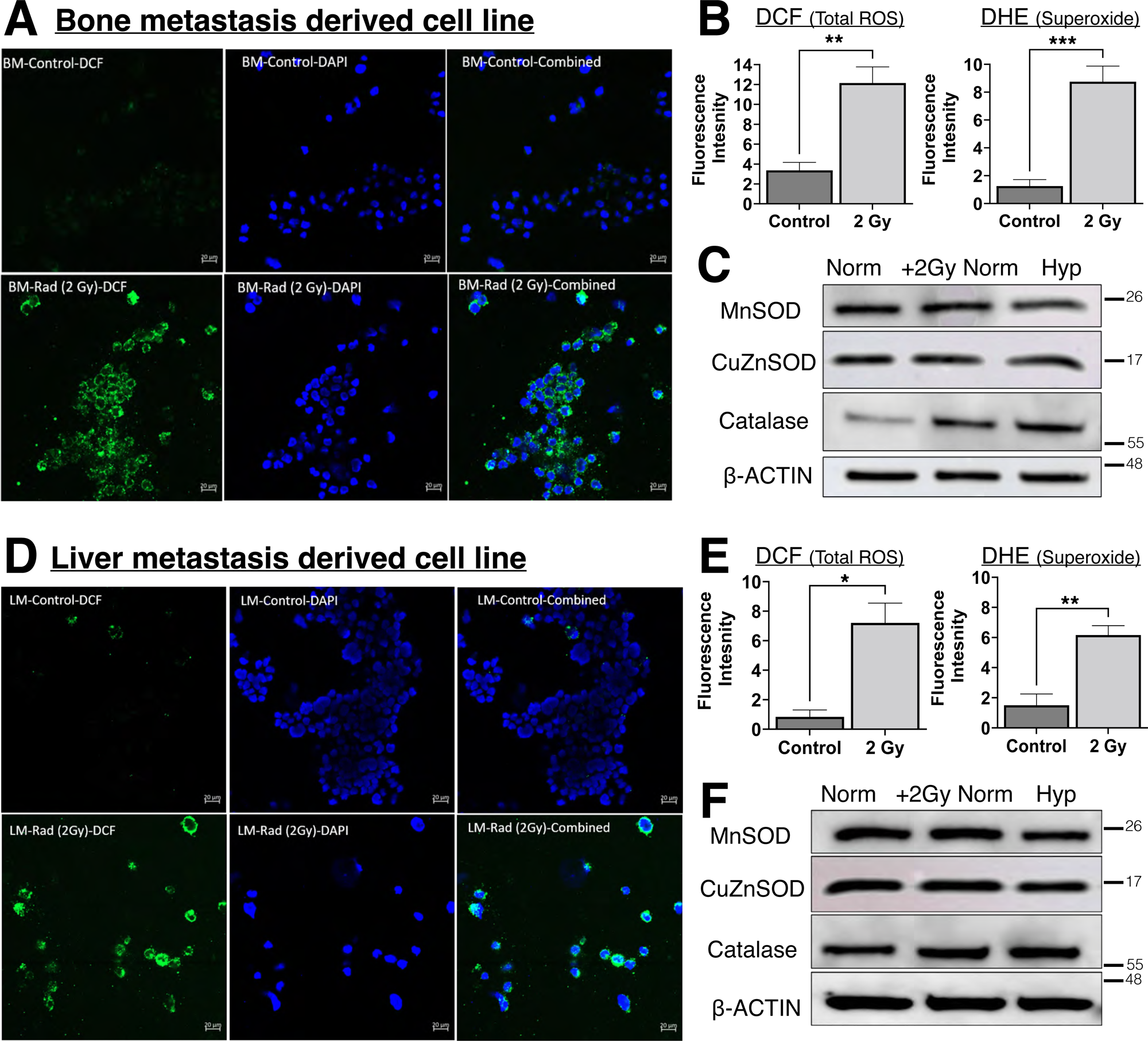
Radiation and hypoxia induce ROS accumulation and an elevated antioxidant defense program in PC RAP-derived cell lines. (A) Representative fluorescence microscopy images of bone metastasis-derived cells stained with DCF (total ROS, green) and DAPI (blue) under control and 2 Gy radiation conditions. Scale bars, 20 µm. (B) Quantification of DCF (left) and DHE (right) fluorescence intensity in bone metastasis-derived cells under control, 2 Gy radiation (normoxia), or hypoxia conditions. Data represent mean ± SEM. Statistical significance determined by unpaired two-tailed Student’s t-test; **p<0.01, ***p<0.001. (C) Western blot analysis of antioxidant defense enzymes MnSOD, CuZnSOD, and Catalase in bone metastasis-derived cells under normoxia (Norm), 2 Gy radiation (+2Gy Norm), or hypoxia (Hyp) conditions. Beta-actin served as loading control. (D) Representative fluorescence microscopy images of liver metastasis-derived cells stained with DCF and DAPI under control and 2 Gy radiation conditions. Scale bars, 20 µm. (E) Quantification of DCF (left) and DHE (right) fluorescence intensity in liver metastasis-derived cells. Data represent mean ± SEM. Statistical significance determined by unpaired two-tailed Student’s t-test; *p<0.05, **p<0.01. (F) Western blot analysis of MnSOD, CuZnSOD, and Catalase in liver metastasis-derived cells under normoxia (Norm), 2 Gy radiation (+2Gy Norm), or hypoxia (Hyp) conditions. Beta-actin served as loading control. MnSOD, manganese superoxide dismutase, CuZnSOD, copper-zinc superoxide dismutase; DCF, 2′,7′-dichlorofluorescein diacetate; DHE, dihydroethidium.

### Modeling the NEPC Bone Metastasis Microenvironment Using a Novel Organoid Co-culture System

A fundamental limitation of conventional organoid monoculture is the absence of the native tumor microenvironment. In the case of bone metastatic CRPC, the complex cellular niche comprises hematopoietic, stromal, endothelial, and immune cell populations that profoundly shape tumor behavior and therapeutic response. Modeling this niche in vitro has historically been hampered by the lack of systems that faithfully recapitulate the cellular composition and 3D architecture of human bone marrow [94]. To address this limitation, we developed a first-of-its-kind co-culture model combining PDO_BMO with iPSC-derived human bone marrow organoids, generated via the stepwise directed differentiation protocol published by Lam et al. [95]. This protocol differentiates human iPSCs to mesenchymal, endothelial, and hematopoietic lineages, producing 3D structures containing stroma, lumen forming sinusoids, and myeloid cells that closely recapitulate the human bone marrow microenvironment [94–98]. Once the iPSC derived human bone marrow organoid had been established, PDO_BMO was added to the culture system. Histopathological evaluation of the co-culture revealed two morphologically distinct cell populations coexisting within a single organoid structure (Figure 9A). The iPSC derived bone marrow organoid component displayed loose, hypocellular architecture with scattered hematopoietic-like cells embedded in a stromal matrix consistent with the marrow-like morphology of the bone marrow organoid cultured alone (Figure 9B). In contrast, PDO_BMO cells formed discrete, tightly packed “tumor nodules” with high-grade morphology, large nuclei, prominent nucleoli, and clearly delineated from the surrounding marrow stroma by a well-defined tumor-stroma interface (Figure 9A, dotted line around PDO_BMO). This architectural organization reflects the histological pattern of CRPC-NE bone marrow infiltration observed in our patient autopsy tissue (Figure 4C), suggesting that the co-culture system can recapitulate key structural features of the in vivo bone metastasis niche. To confirm the identity of each cellular compartment within the co-culture, we performed multiplex IF staining for KRT8/18 and VIM (Figure 9C). The luminal epithelial marker KRT8/18 was retained in PDO_BMO, strongly and specifically labeling the organoids. Whereas VIM, a mesenchymal marker expressed by stromal and hematopoietic cells, was expressed in the surrounding bone marrow organoid compartment. DAPI nuclear counterstaining confirmed the distinct spatial organization of the two populations. The compartmentalization of KRT8/18 positive tumor cells and VIM positive marrow stroma demonstrates that PDO_BMO maintains its epithelial tumor identity within the bone marrow niche. The two populations of cells organize spatially in a manner reminiscent of tumor-stroma interactions observed in vivo. Notably, the co-culture revealed scattered cells (high-magnification H&E; Figure 9A) with high-grade morphology within the iPSC bone marrow organoid compartment, and KRT8/18 positive cells were observed intermixed within the VIM positive marrow stroma on IF. This suggests active infiltration of PDO_BMO cancer cells into the bone marrow organoid niche. While further studies are needed to confirm whether this represents directed invasion or passive mixing during co-culture establishment, this observation raises the intriguing possibility that PDO_BMO cells retain an intrinsic capacity to infiltrate and colonize a human bone marrow-like microenvironment in vitro, thereby recapitulating a key step in the metastatic cascade. This co-culture system represents a first step toward modeling the bone metastasis tumor microenvironment of CRPC-NE in vitro, and establishes an innovative platform for future mechanistic studies of tumor-niche crosstalk and the development of bone metastasis directed treatment strategies [94, 97].

**Figure 9.**
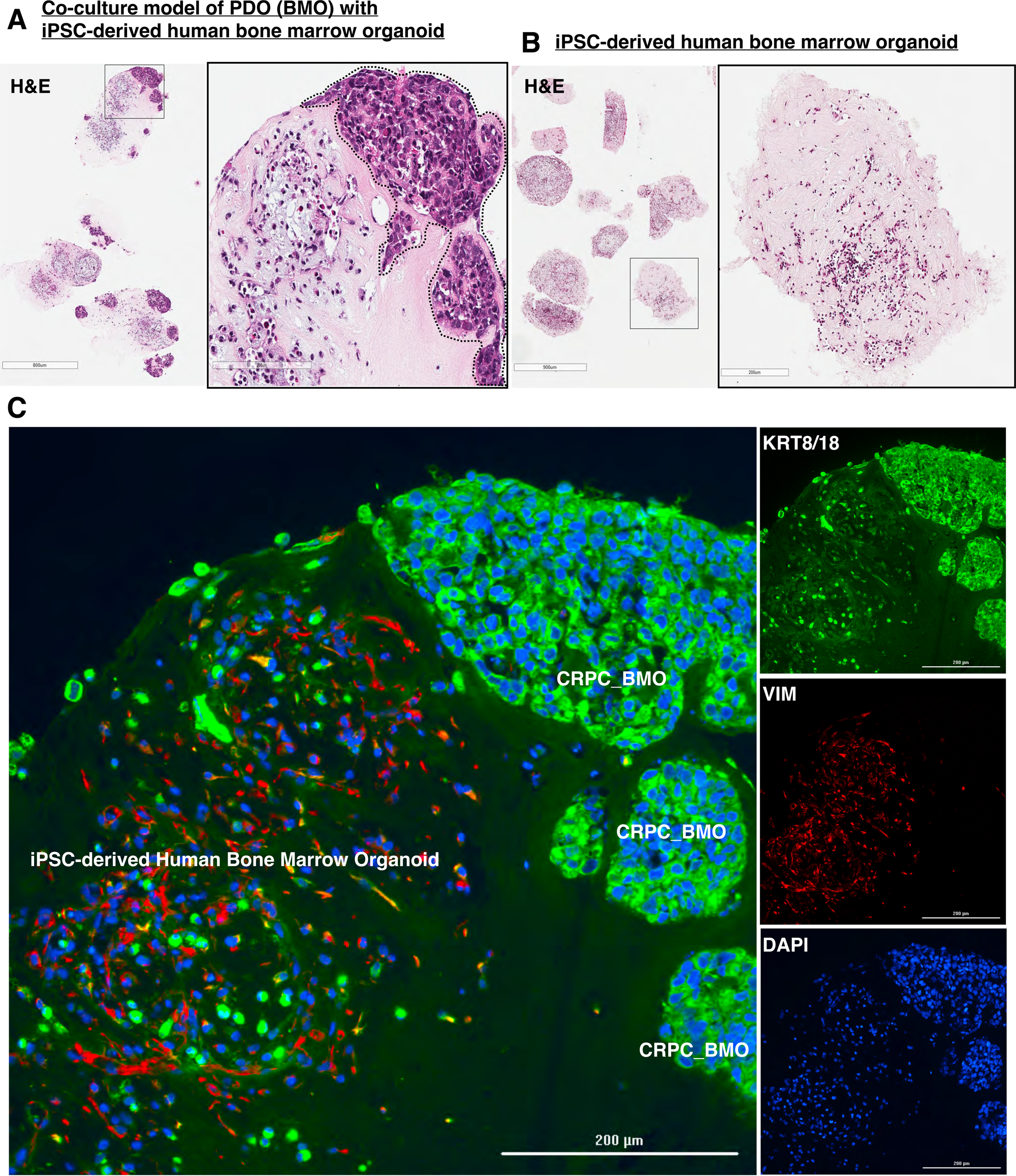
Co-culture of PDO_BMO with iPSC-derived human bone marrow organoid models the bone metastasis microenvironment. (A) H&E staining of the PDO_BMO and iPSC-derived human bone marrow organoid co-culture at low magnification (left; scale bars, 800 µm) and high magnification (right; scale bars, 200 µm). The dotted line delineates the interface between the PDO_BMO tumor nodule and the surrounding iPSC-derived bone marrow organoid stroma. (B) H&E staining of the iPSC-derived human bone marrow organoid alone at low magnification (left; scale bars, 900 µm) and high magnification (right; scale bars, 200 µm), demonstrating loose marrow-like architecture with scattered hematopoietic cells embedded in stromal matrix. (C) Multiplex immunofluorescence of the co-culture showing KRT8/18 (green, epithelial tumor marker), Vimentin (VIM, red, mesenchymal/ stromal marker), and DAPI nuclear counterstain (blue) at low magnification (left; scale bars, 200 µm) and high magnification individual channels (right; scale bars, 200 µm). PDO_BMO tumor nodules are labeled.

### RAP-Derived PDOs Establish Spontaneously Metastatic In Vivo Models of Lethal NEPC

To build in vivo preclinical models from PC RAP-derived tissue, we attempted to establish patient-derived xenografts (PDX) and patient-derived organoid xenografts (PDOX) models. Using publicly available transcriptomic datasets generated by the NCI PDMR from the donor patient, we assessed transcriptomic concordance with our independently established organoid lines. This powerful analysis was a cross-institutional, multi-dataset validation of our PDO models. To enable direct comparison, we re-analyzed RNA-seq FASTQ files from the two NCI PDMR source patient tumors and eleven patient-matched PDX models using the same pipeline applied to our PDOs. We performed a Variance Stabilizing Transformation to normalize expression values for a curated NEPC gene panel, which was used to generate NEPC and adenocarcinoma (Adeno) scores for all samples [6]. The NEPC score applied in our analysis is the 70 gene integrated classifier developed and validated by Beltran et al. [6]. This standard transcriptomic benchmark for NEPC molecular classification was originally shown to distinguish CRPC-NE from CRPC-Adeno [6]. Unsupervised hierarchical clustering of these samples, along with our PDO lines and LNCaP adenocarcinoma controls, revealed three clearly distinct groups (Figure 10). LNCaP cells, representing canonical AR driven adenocarcinoma, clustered separately from all other samples, with high Adeno scores and low NEPC scores, confirming the expected lineage separation. In contrast, PDO_BMO and PDO_LMO clustered with the PDMR originator patient tissue samples (ORIGINATOR_331_R2 and R4) and all eleven PDX models (NCI PDMR repository) derived from the same liver metastasis, sharing uniformly high NEPC scores and low Adeno scores throughout. Both originator samples demonstrated high NEPC scores (0.62) and low Adeno scores (0.13 and 0.15), consistent with their clinical designation as treatment emergent NEPC. The PDX samples (NCI PDMR repository) showed NEPC scores comparable to our PC RAP derived PDO lines, confirming transcriptomic fidelity across multiple rounds of passaging the PDOs (passage 10). At the gene level, the NEPC cluster, comprising PDO_BMO, PDO_LMO, originator tissue, and PDX models, was characterized by high expression of canonical NEPC signature genes (Figure 10, asterisk), including *EZH2*, *AURKA*, *SYT11*, *PROX1*, *MYCN*, and *SCG3*, alongside marked downregulation of *AR* driven luminal genes, including *AR*, *KLK3*, *KLK4*, *NKX3-1*, and *HOXB13* [6, 24]. Notably, RB1 expression was reduced across the NEPC cluster relative to LNCaP controls, consistent with the genomic LOH at the RB1 locus identified by WES analysis (Figure 3) and the protein level reduction observed by western blot (Figure 6C). Taken together, these data provide robust cross-institutional transcriptomic validation of our PC RAP derived organoid lines, demonstrating that PDO_BMO and PDO_LMO recapitulate the molecular identity of the source patient tumor validated across multiple independent model systems and institutions [6, 24].

**Figure 10.**
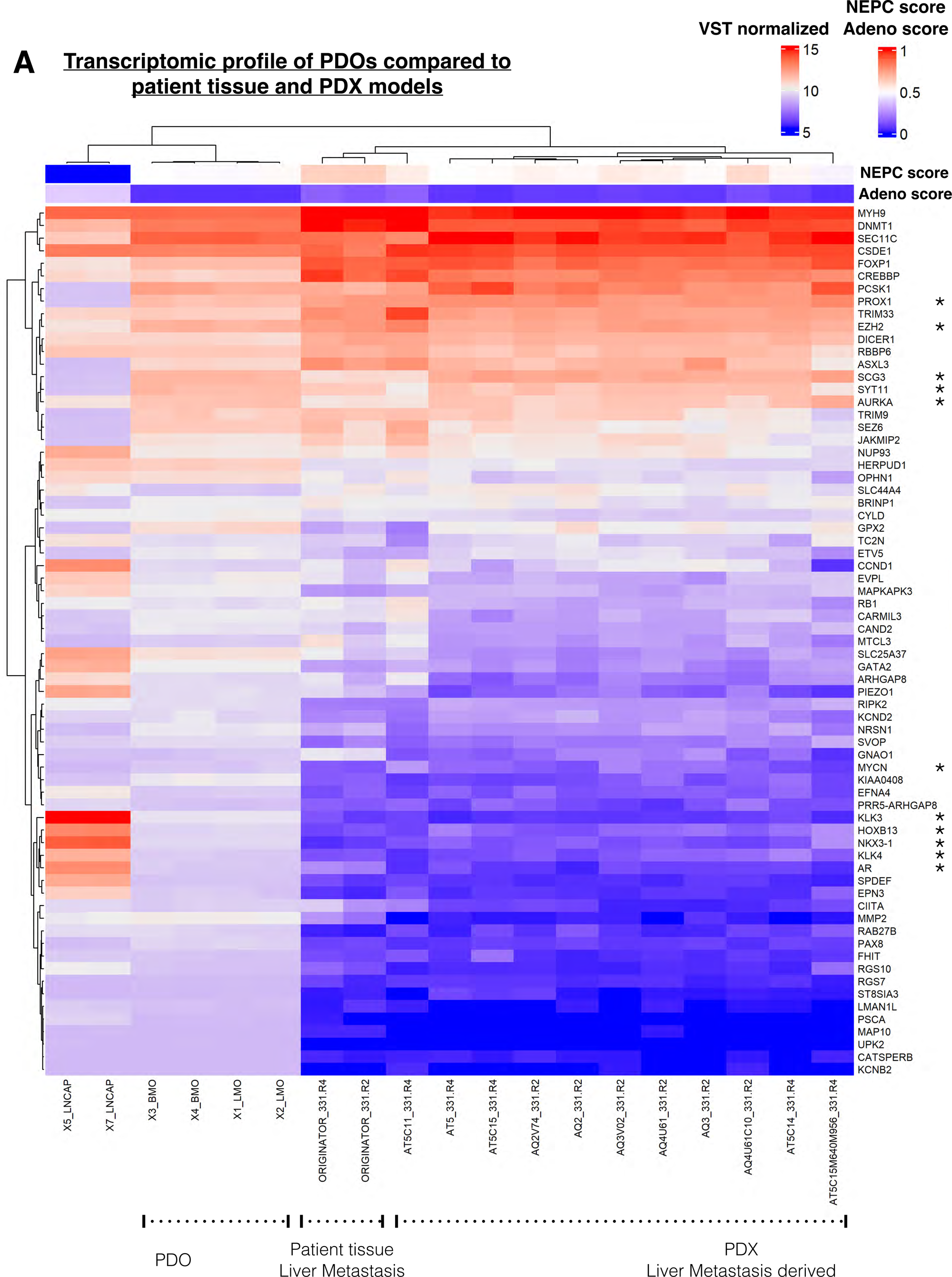
Transcriptomic profiling of PC RAP-derived organoids compared to patient tissue and PDX models demonstrates cross-institutional concordance. (A) Unsupervised hierarchical clustering heatmap of VST normalized RNA sequencing expression values for a curated NEPC and adenocarcinoma gene panel across three sample groups: PC RAP-derived PDO lines (PDO_BMO and PDO_LMO) and the adenocarcinoma cell line LNCaP as control (left); source patient liver metastasis tissue samples (ORIGINATOR_331_R2 and ORIGINATOR_331_R4) (center); and eleven liver metastasis-derived PDX samples from the NCI Patient-derived Models Repository (PDMR; right). NEPC (top bar) and adenocarcinoma (Adeno) scores are represented as a red-blue gradient (bottom bar); individual scores are shown for each sample. Gene expression values are shown as VST-normalized counts (color scale: blue, low; red, high). RNA-seq FASTQ files from PDMR originator and PDX samples were re-analyzed using the same bioinformatics pipeline applied to PC RAP-derived PDO samples.

Having established the molecular identity of our PC RAP derived PDO lines, we next sought to evaluate their tumorigenic potential in vivo through the generation of PDX and PDOX models. While the bone metastasis derived tissue did not engraft successfully, the liver metastasis derived PDX was successfully established and was labeled as PDX_LM. The tumors from PDX_LM were serially propagated through passage 3. Histopathological analysis of PDX_LM confirmed preservation of the mixed neuroendocrine and adenocarcinoma phenotype of the source tumor, with H&E demonstrating high-grade carcinoma morphology consistent with the source patient tissue (Figure 11A). IF staining of passage 3 PDX_LM tumors revealed KRT8/18 and NCAM1 expression, along with a high Ki-67 proliferative index, in the PDX_LM (Figure 11A). The PDX tumors maintained expression of SYP, confirming preservation of the NEPC phenotype through successive in vivo propagation (Supplementary Figure 65A). EZH2 expression was similarly retained, confirming maintenance of the epigenetic reprogramming signature characteristic of the originating tissue and supporting the role of PRC2 mediated chromatin remodeling in sustaining the NEPC phenotype in vivo (Supplementary Figure 5A). During routine PDX propagation, we observed spontaneous lymph node (LN) metastasis in 1 out of 5 tumor bearing mice. IF characterization of the metastatic LN lesions demonstrated preserved expression of NEPC markers, including KRT8/18, NCAM1, and Ki-67 (Supplementary Figure 5B), confirming that PDX_LM retains metastatic capacity in vivo and maintains the neuroendocrine differentiation phenotype in disseminated disease. Co-immunostaining of PDX_LM tumor demonstrated loss of basal cytokeratins (KRT5/14) with strong retention of luminal cytokeratins (KRT8/18), consistent with luminal NEPC differentiation (Supplementary Figure 6A). Collectively, these findings validate the tumorigenic and metastatic potential of PDX_LM and confirm its phenotypic resemblance with the source patient tumor [12, 13].

**Figure 11.**
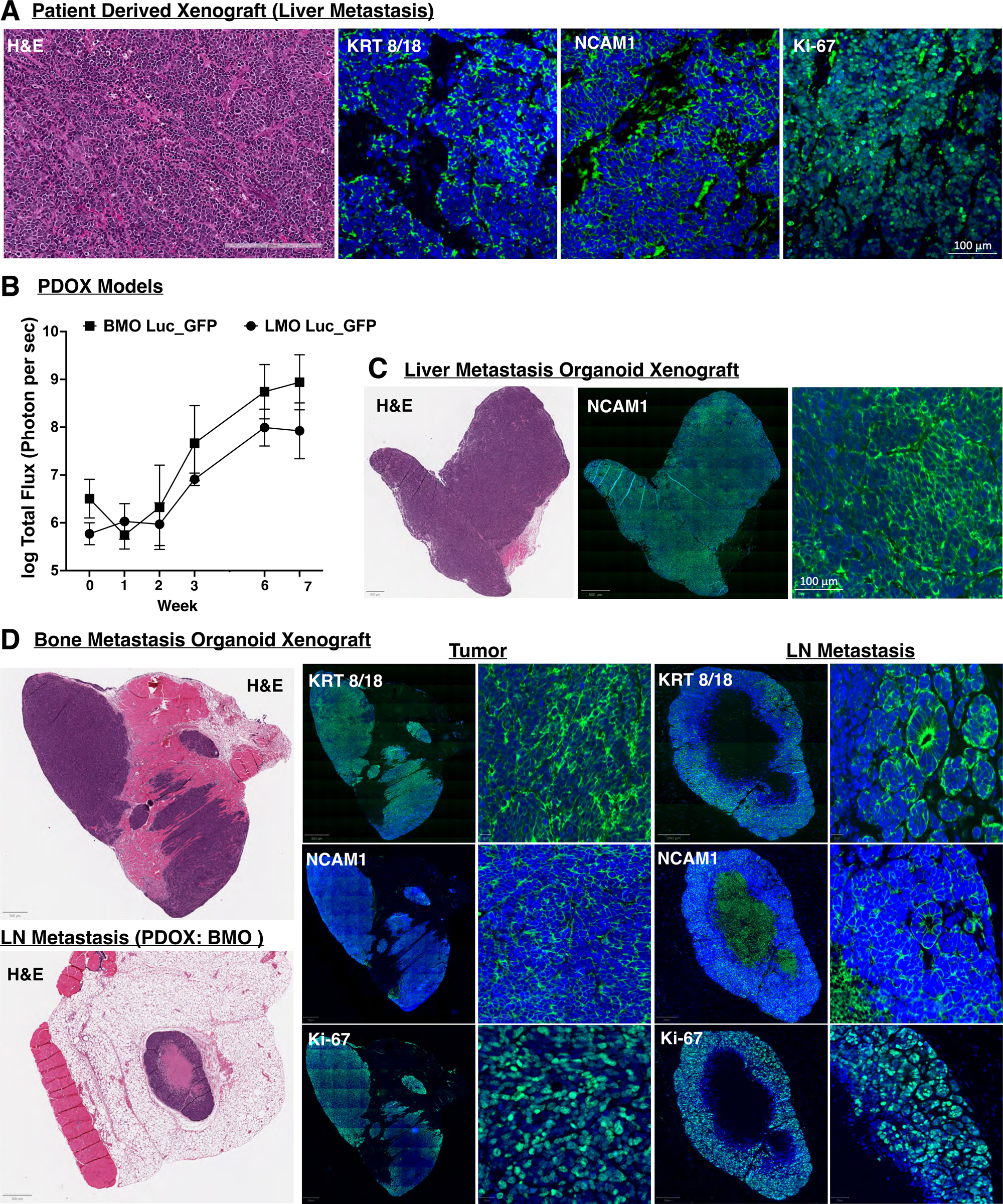
PC RAP-derived organoid lines establish tumors and recapitulate metastatic behavior in vivo. (A) Histopathological and immunofluorescence characterization of the liver metastasis patient-derived xenograft (PDX), showing H&E staining (scale bars, 200 µm) and multiplex immunofluorescence for KRT 8/18, NCAM1, and Ki-67 (green) with DAPI nuclear counterstain (blue). Scale bars, 100 µm. (B) In vivo tumor growth curves of luciferase and GFP expressing PDO_BMO (squares) and PDO_LMO (circles). PDOX models were measured by bioluminescence imaging over seven weeks. Data represent log total flux (photons per second) ± SEM; n=4 mice per group. (C) Histopathological and immunofluorescence characterization of the LMO PDOX, showing H&E (scale bars, 500 µm) at low magnification and NCAM1 immunofluorescence at low (scale bars, 800 µm) and high magnification (scale bars, 100 µm). (D) Histopathological and immunofluorescence characterization of the BMO PDOX primary tumor and spontaneous lymph node (LN) metastasis, showing H&E staining (scale bars, 500 µm) and immunofluorescence for KRT 8/18, NCAM1, and Ki-67 (green) with DAPI nuclear counterstain (blue) at low (scale bar, 800 µm) and high magnification (scale bar, 20 µm) in both compartments.

Since direct PDX establishment from the PC RAP derived bone metastasis tissue was unsuccessful, we asked whether our patient-derived organoid lines (LMO and BMO) could serve as an alternative in vivo preclinical platform. To address this, we generated PDOX models by implanting luciferase/GFP-expressing PDO_BMO and PDO_LMO into immunocompromised mice and monitored tumor growth by bioluminescence imaging over seven weeks. Both lines demonstrated progressive tumor establishment and growth, with log total flux increasing consistently from baseline through week 7 (Figure 11B). PDOX_BMO showed robust, slightly faster tumor growth than PDOX_LMO, though both lines reliably established tumors, confirming the tumorigenic capacity of PC RAP-derived organoid lines in vivo. Histopathological and IF characterization of both PDOX models confirmed retention of the NEPC molecular phenotype in vivo, with robust NCAM1 expression (Figures 11C and D) and a luminal differentiation profile (KRT5/14 loss, KRT8/18 retention; Figures 11C and D; Supplementary Figure 6) similar to the parental PDOs, PDX models, and source metastases (Figure 7 and Supplementary Figure 5A). H&E staining revealed high-grade carcinoma morphology in both models, with strong Ki-67 proliferative activity. Notably, PDOX_BMO produced spontaneous lymph node metastases that preserved KRT8/18, NCAM1, and Ki-67 expression (Figure 11D), demonstrating retention of the metastatic and invasive capacity of the source bone metastasis [30, 33].

## DISCUSSION

In this study, we report the establishment and comprehensive characterization of novel patient-derived preclinical models of bone and liver metastases from treatment emergent PC. By establishing cellular models from two anatomically and biologically distinct metastatic sites, we provide a unique preclinical resource for studying the biology and therapeutic vulnerabilities of lethal PC. Using a standardized tissue processing workflow, we have successfully generated organoid lines (PDO_LMO and PDO_BMO), an MSC cell line, PDX, and PDOX models from a single donor enrolled in the UNMC RAP. Both PDO lines have been stably propagated beyond passage 10 and have faithfully preserved the molecular and histopathological features of the source tumor across multiple independent platforms and institutions.

The scientific output of our study highlights the infrastructure and capabilities of the PC-RAP. First and foremost is truly the ultimate gift given by patients, based on their trust of the clinicians who care for them. The success of any RAP depends on the extraordinary generosity of patients and their families, who make the deeply personal decision to donate tissue for research [99]. The patient from whom these models were derived survived two years from diagnosis, but chose to spend that time as both a patient and a scientific partner. The preclinical models generated in this manuscript are a reflection of that choice, and honoring it through rigorous, impactful science is our most important obligation. This volunteer driven, multidisciplinary effort demonstrates how team science among clinicians, pathologists, and basic scientists can sustain a logistically demanding research infrastructure. Importantly, participation in the RAP provides a unique training opportunity for graduate students, residents, and postdoctoral fellows, exposing them to the real-world consequences of the diseases they study in the laboratory. Programs like our PC RAP generate scientific resources while training the next generation of researchers. We further demonstrate how collaborative team science enabled multiple laboratories to combine protocols and resources from a single patient’s autopsy to generate a diversity of models for studying lethal PC. Establishing a successful RAP requires substantial long-term investment in planning, funding, and institutional support [100]. Our study demonstrates that multi-laboratory partnerships maximize the scientific value of each donation and should be adopted as a standard practice.

The donor patient’s clinical trajectory represented the aggressive phenotype of treatment emergent NEPC. Following diagnosis with high grade prostate adenocarcinoma (Gleason group 5) and subsequent ADT, the patient developed castration resistance and histological transformation to small cell CRPC-NE within 14 months. This timeline is considerably shorter than the median interval of 39.7 months reported between adenocarcinoma diagnosis and treatment emergent NEPC [30]. Moreover, the patient’s treatment course comprised of a multi-agent, non-targeted approach, highlighting the lack of approved NEPC specific therapies [33]. At autopsy, widespread metastatic disease with mixed adenocarcinoma and NEPC histology was observed, a pattern seen in 40-50% of NEPC cases and associated with modestly improved survival compared to pure small cell carcinoma, though outcomes remain uniformly poor [30].

Despite its recognized importance, NEPC models remain rare. To our knowledge, the PDOs generated in this study represent among the few preclinical models of bone and visceral metastases derived from NEPC tissue. The NCI-H660 cell line, a commercially available patient-derived NEPC cell line, grows slowly as floating clusters, making it technically challenging for routine in-vitro assays. Our PDO lines grow in both 2D and 3D culture, offering a tractable and biologically relevant platform to study dependencies in treatment emergent PC. Transcriptomic comparison with NCI-H660 demonstrated close clustering by PCA, confirming a shared NEPC molecular identity with our models. NCI-H660 was derived from a lymph node metastasis initially diagnosed as small cell lung carcinoma [8] and subsequently reclassified as prostate in origin [9]. It is therefore thought to represent a de novo NEPC model, distinct from treatment emergent NEPC, which arises on an adenocarcinoma background [71]. In contrast, our PDO lines are derived from a patient who progressed through adenocarcinoma to treatment emergent NEPC under the therapeutic pressure of ADT. This distinction is biologically important and relevant from a clinical and translational perspective.

The comprehensive molecular characterization of the PDO lines using orthogonal techniques represents one of the strongest aspects of this study. Together, these analyses provide complementary lines of evidence supporting molecular similarity to the patient tumor. At the genomic level, analysis of publicly available NCI PDMR whole exome sequencing data from the originator liver metastases revealed a mutational landscape highly consistent with treatment emergent NEPC [6, 29]. We identified a clonal gain-of-function hotspot mutation in *TP53* (p.R273H, VAF 0.95-0.99), a loss-of-function mutation in *ARID1A*, *BRCA1* missense mutation, and *ATM* mutation. In addition, we detected copy-neutral LOH at the *PTEN* locus on chromosome 10 and at the *RB1* and *BRCA2* loci on chromosome 13. Notably, the *TP53* p.R273H mutation is one of the most frequently occurring hotspot mutations in human cancer, classified as a gain-of-function alteration that not only abrogates the tumor suppressive activity of wild-type p53 but also confers oncogenic properties, including enhanced invasion and therapy resistance [101–106]. In PC, co-alternations in the tumor suppressors *PTEN*, *TP53,* and *RB1* cooperate mechanistically to drive lineage plasticity, neuroendocrine transdifferentiation, and metastatic progression [107–111]. Cumulative alterations in these genes are associated with poor prognosis, faster disease progression, and reduced overall survival. Patient genomics and mouse genetic models have further demonstrated that these co-deletions promote metastasis and therapeutic resistance [55, 112–115], while also revealing targetable vulnerabilities and key downstream mediators such as EZH2 and SOX2 [74, 116, 117]. Our PDOs recapitulate this genetic context and express *EZH2* and *SOX2*, functional drivers of lineage plasticity and treatment resistance [74, 75, 116, 118]. Importantly, prior studies have demonstrated that *SOX2* knockdown in the context of *PTEN/RB1/TP53* loss restores enzalutamide sensitivity and downregulates NE associated gene expression [74, 75]. Building on this, work from the Rizzino laboratory has shown that elevating *SOX2* in PC cells directly upregulates the expression of a broad panel of neuroendocrine genes, including synaptophysin, while also suppressing canonical AR driven luminal gene networks [85, 86].

The cross-institutional transcriptomic validation performed in this study provides strong evidence that our PDO models resemble their tissue of origin. Using publicly available RNA-seq data from the NCI PDMR, we show that PDO_LMO and PDO_BMO (passage 10) cluster tightly with the originator tissue and eleven matched PDX models across independent datasets and institutions. All samples exhibited high neuroendocrine and low adenocarcinoma scores. This analysis confirms that the NE transcriptomic identity is not only preserved in our models but is stable through serial propagation (NCI PDMR). Our cross-dataset analysis of independently generated model systems across two institutions UNMC and the NCI PDMR, is particularly compelling as it strengthens the translational credibility of our models. Furthermore, the PDOs generated in this study exhibit a protein expression profile consistent with treatment emergent NEPC, including loss of AR, NKX3.1, and PTEN, and expression of neuroendocrine/stemness markers. They also retain an intermediate phenotype with high EZH2 and SOX2 expression, indicating epigenetic lineage plasticity.

A unique and novel aspect of this study is the characterization of mesenchymal stem cells (MSCs) isolated from bone fragments adjacent to the spinal metastasis. These findings demonstrate successful extraction of a stromal cell population from a PC patient’s bone metastases. Based on their expression of classical phenotypically MSC markers, the absence of hematopoietic cell markers, and capacity to differentiate into adipocytes, these cells are likely to be MSCs. While they lack the osteogenic differentiation capacity characteristic of MSCs [60], there are several factors that may explain these patient-derived MSCs’ inability to differentiate into osteoid-producing cells/osteoblasts. It is important to consider the impact of the metastatic tumor cells on adjacent MSCs. Although numerous works provide evidence for altered MSC function in the tumor microenvironment, there have been no studies, to date, to isolate and characterize MSCs from PC patient spinal metastases i.e., MSCs adjacent to metastatic prostate tumors in bone [119–121]. Specifically, RAP MSCs were derived from small cell neuroendocrine PC, a subtype which commonly induces osteolytic rather than osteoblastic bone metastases [39, 122–125]. Therefore, the inability of RAP MSCs to produce osteoid may be indicative of an osteolysis-promoting phenotype of bone stromal cells in small cell carcinoma metastases. This was supported by increased RANKL expression in comparison to healthy MSCs however, further study is needed to determine the functional impact of co-upregulation of RANKL and OPG, as osteoclast formation depends upon the ratio of these two factors [126]. Additionally, RAP MSC ability to differentiation into adipocytes may indicate tumor-mediated skewing towards that pathway since adipocytes have been shown to promote BM-PC progression through multiple mechanisms, including promotion of proliferation, migration and invasion, oxidative and ER stress resistance, and tumor-induced osteolysis [127–132]. The phenotype of the RAP MSCs may be a result of other patient-specific factors. Sex, ethnicity, age, lifestyle, environmental factors, and other medical conditions could all contribute to the differences observed between these MSCs and those from healthy donors. Age in particular is a likely factor in their lack of osteogenic differentiation, based on prior studies showing decreased osteogenic potential of MSCs from aged versus young mice [133]. These patient-derived MSCs also expressed lower levels of CXCL12/CXCR4, which several studies have linked to decreased osteogenic potential of aged mouse MSCs [133]. For future studies, isolation of MSCs from distant bone regions for comparison to metastasis-adjacent MSCs could help account for inter-patient and therapy induced variation and enhance analyses of tumor-induced changes in patient-derived MSCs. To our knowledge, this is the first characterization of MSCs isolated from a NEPC spinal metastasis, establishing a novel ex vivo cellular model for studying the bone-tumor interface. Together with our PDO lines, this resource enables investigation of tumor-stromal crosstalk in the context of NEPC bone metastasis. Similarly, the proof-of-concept PDO and iPSC derived bone marrow organoid co-culture system further extends our modeling capacity toward the tumor microenvironment. To model the bone metastatic niche, we established a co-culture system combining the bone metastasis derived PDO (PDO_BMO) with iPSC-derived bone marrow organoid system (published differentiation protocols [94, 95, 98]), comprising endothelial, stromal, and hematopoietic cell populations. We observed PDO integrating within the iPSC bone marrow organoids, providing a novel, scalable platform to study cellular interactions in the bone metastatic niche.

Elevated antioxidant defense as a mechanism of radioresistance has been reported in multiple cancer types. It is well documented in cancer stem cell populations, where high ROS scavenging capacity is thought to confer resistance to oxidative damage from both radiation and chemotherapy [92]. Our findings suggest that both PDO lines can tolerate elevated ROS under radiation and hypoxia conditions. In the NEPC context, the antioxidant defense pathway represents a potential therapeutic vulnerability, as pharmacological inhibition of ROS scavenging enzymes using agents such as auranofin (thioredoxin reductase inhibitor) or buthionine sulfoximine (glutathione synthesis inhibitor) has shown pre-clinical efficacy in sensitizing radioresistant cancer cells to ionizing radiation [134]. The cell lines generated in this study are an ideal system for evaluating such combinatorial strategies in the NEPC context. Future studies incorporating clonogenic survival assays and irradiation of PDX/ PDOX models will be essential to fully characterize the potential radioresistance phenotype and its therapeutic implications.

While conventional PDX models from patient tissue have low engraftment rates, PDOX models derived from organoid lines achieve nearly 100% engraftment rates. This approach substantially improves upon the historically low success rates of direct PDX establishment from CRPC samples [13, 135]. PDOX models are fairly easy to generate, scalable, and recapitulate the in vivo complexity of xenograft biology [135]. Of particular relevance to our study is the successful establishment of PDOs and subsequent PDOX models from bone metastases. This is an important methodological advance, given the difficulty in generating conventional PDX models from bone. Our complementary approach therefore addresses a fundamental limitation of standard PDX workflows and broadens the range of clinically relevant disease sites that can be modelled experimentally. Furthermore, our PDOX models retain the histological and molecular features of the parent organoids and source tissue, can be serially propagated, and provide a platform for studying tumor biology *in vivo*. Other applications include metastasis modeling, tumor stroma interactions, vascular recruitment, and in vivo drug pharmacokinetics. The spontaneous lymph node metastasis observed in our PDOX models demonstrates the unique capacity of PDOX systems to reveal functional biological differences between models. These PDOX models support preclinical studies of drugs targeting NEPC drivers (EZH2, SOX2), PARP inhibitors in BRCA1 mutant backgrounds, synthetic lethality strategies for *PTEN/RB1/TP53* null PC, and antioxidant inhibitors as radiosensitizers. Moreover, the ability to move from organoid drug screening to in vivo PDOX validation within a single patient-derived system preserves genetic/tumor characteristics while accelerating precision therapy development for NEPC [13, 24].

Collectively, this work provides a blueprint for utilizing rapid autopsy tissue to generate valuable preclinical models. The program is sustained by a committed multidisciplinary volunteer team and the altruism of patients who consent to postmortem tissue donation. In this study, we have established novel preclinical models consisting of patient-derived organoids, MSC cultures, PDX, and PDOX models. These models faithfully recapitulate the molecular and functional characteristics of the source patient tumor across multiple independent platforms and institutions. The models exhibit intrinsic radioresistance, can be integrated into co-culture systems to model the metastatic niche and demonstrate spontaneous metastatic capacity in vivo. Together, they constitute a biologically relevant platform for interrogating the biology and accelerating the development of effective therapeutic strategies for patients with lethal PC.

### Limitations of the study

We acknowledge that there are several limitations of this study. First and most importantly, our PDO lines are derived from a single patient. While the clinical trajectory and molecular phenotype of this patient are representative of aggressive treatment emergent NEPC, single patient models inevitably capture only a snapshot of the biological and genomic diversity of the disease. The extent to which the specific dependencies and drug response profiles of the PDO lines match the broader NEPC patient population remains to be determined. Expanding the PDO biobank across multiple PC patients will be essential to address this question. Second, organoid culture by definition removes tumor cells from their native microenvironment. Tissue microenvironment is a critical mediator of tumor cell behavior, therapeutic response, and metastatic biology in vivo. Although we have addressed this limitation through the co-culture of PDO_BMO with iPSC derived human bone marrow organoids, this system does not yet incorporate the full complexity of the bone metastasis niche. These include immune cell populations, osteoclasts, osteoblasts, and tumor vasculature components that are increasingly recognized as critical determinants of NEPC progression and therapy resistance. Third, the response to radiation/hypoxia data presented here are proof-of-concept observations from 2D cell cultures under defined radiation and hypoxia conditions, and do not yet reflect the complexity of the in vivo tumor microenvironment or clinical radiation regimen. Validation of radioresistance mechanisms and the therapeutic relevance of the antioxidant defense program highlighted in our study will require additional functional studies, including clonogenic survival assays, in vivo irradiation models, and correlation with clinical response data. Fourth, while our PDOX models demonstrate successful in vivo tumor establishment and spontaneous lymph node metastasis, the immunocompromised host used for xenograft studies precludes assessment of the tumor-immune cell interactions. Finally, while the PC RAP workflow is powerful, it is logistically demanding and dependent on sustained engagement from a multidisciplinary volunteer team. An efficient, well-coordinated team is critical for minimizing variability in tissue quality, which can be influenced by both the post-mortem interval and the anatomical site of collection. Overall, addressing these limitations through expansion of the PC RAP PDO biobank, integration of matched multi-omic profiling, development of more complex co-culture and humanized mouse systems, and collaboration across PC RAP programs at other institutions will be essential next steps to maximize the translational impact of these models.

## RESOURCE AVAILABILITY

## Lead contact

## Materials availability

Tissue specimens were deposited at the NCI PDMR, NCI-Frederick, Frederick National Laboratory for Cancer Research, Frederick, MD. The PDX models described in this study are available upon request through the NCI PDMR (https://pdmr.cancer.gov/) or the UNMC RAP, subject to applicable material transfer agreements and institutional approvals.

## Data and code availability

Raw RNA-seq data generated from patient-derived organoids have been deposited in the NCBI Gene Expression Omnibus (GEO; GSE) and will be released upon publication. Raw sequencing data for WGS and RNA-seq of RAP patient-derived PDX models were downloaded from the NCI PDMR. URL: https://dctd.cancer.gov/drug-discovery-development/reagents-materials/pdmr. Raw data may be requested from with appropriate institutional approvals.

The software and algorithms for data analyses used in this study are published and referenced throughout the methods section.

This study does not report original code.

Any additional information required to reanalyze the data reported in this paper is available from the lead contact upon request.

## ACKNOWLEDGEMENTS

This work was supported by the National Institutes of Health/National Cancer Institute, NIGMS COBRE (P20GM121316), R01CA274605, American Cancer Society Institutional Research Grant, and FPBCC pilot grants to G. Mathew, and the APC was funded by the same awards. R.C. Bergan R. Rhatigan were supported by NCI R01CA276846. M. Abdalla and R. Salloom were supported by an American Cancer Society Research Scholar Grant (RSG-) and FPBCC pilot grants. T.C. Caffrey, P.M. Grandgenett and M.A. Hollingsworth were supported by NCI Cancer Center Support Grant, P30CA36727. P.M. Grandgenett was additionally supported by the NCI Research Specialist, R50CA211462. L.M. Cook and C.S. Johnson were supported by an American Cancer Society Research Scholar Grant (RSG-19-127-01-CSM) and NCI R01CA274605. C.S. Johnson was additionally supported by funding from the NCI and T32 Cancer Biology Training Grant (T32CA009476). G.Ghosal was supported by the NCI 1R01CA263504-01.

The authors thank all members of the RAP volunteer program, pathologists, and study coordinators. Members of the Bergan, Abdalla, Cook, and Mathew laboratories for helpful discussions and critical reading of the manuscript. We are grateful to the Meng laboratory for access to bioprinting and organotypic platforms. We acknowledge the institutional core facilities e.g., genomics, imaging, histology) for technical assistance, and the iCaRe2 biospecimen repository for access to patient-derived specimens. We thank the Live On Nebraska, Project HOPE, and GU Oncology Translational Working Group at the Fred and Pamela Buffett Cancer Center for coordinating tissue donation and the rapid procurement of specimens. We are especially grateful to the patients and their families, whose participation in the Prostate Cancer Rapid Autopsy Program made this work possible.

We acknowledge use of the University of Nebraska Medical Center: 1. UNMC Advanced Microscopy Core Facility, RRID: SCR_022467, P20 GM103427, P30 GM106397, P30 CA036727, S10 RR027301, S10 OD030486, Nebraska Research Initiative. 2. The In Vivo Imaging Spectrum core facility is supported by Cancer Center Support Grant P30 CA036727 and COBRE P20 GM121316. 3. The Organoids core is part of the Nebraska Center for Molecular Target Discovery and Development and supported by the COBRE P20 GM121316. 4. Tissue Sciences Facility, RRID: SCR_012465, which receives support from the Cancer Center Support Grant P30 CA036727 and the UNMC Department of Pathology, Microbiology, and Immunology. Research reported in this publication was also supported by the National Cancer Institute of the National Institutes of Health under award number P30 CA036727 (Cancer Center Support Grant). The content is solely the responsibility of the authors and does not necessarily represent the official views of the National Institutes of Health. The authors used Grammarly to improve the quality of the writing for specific sections of the manuscript.

## AUTHOR CONTRIBUTIONS

Conceptualization, M.A., L.M.C, R.C.B. and G.M.; methodology, B.R., T.C.C, S.S., C.J., R.J.S, K.M., F.M., A.R.B, S.L., P.M.G, M.A., L.M.C, R.C.B. and G.M.; investigations, B.R., T.C.C, S.S., C.J., R.J.S, K.M., A.R.B, P.M.G, M.A., L.M.C, R.C.B. and G.M.; formal analyses, B.R., S.S., C.J., R.J.S, K.M.; data curation, B.R., T.C.C, S.S., C.J., R.J.S, K.M., A.R.B, P.M.G, M.A., L.M.C, R.C.B. and G.M.; resources, Q.P.L, A.R.B., P.M.G., M.A., L.M.C, R.C.B. and G.M.; writing, B.R., C.J., R.J.S, K.M., M.A., L.M.C, R.C.B. and G.M.; funding acquisition, M.A., L.M.C, R.C.B. and G.M.

The contributions (manuscript writing and revisions) of the NIH author (L.M.C) were made as part of their official duties as NIH federal employees, are in compliance with agency policy requirements, and are considered Works of the United States Government. However, the findings and conclusions presented in this paper are those of the authors and do not necessarily reflect the views of the NIH or the U.S. Department of Health and Human Services.

## DECLARATION OF INTERESTS

## EXPERIMENTAL MODEL AND STUDY PARTICIPANT DETAILS

All work involving human tissue and patient-derived material was approved by the UNMC IRB 460-19-EP and 440-16-EP. Informed consent was obtained from all participants or their legal representatives before tissue acquisition through the UNMC RAP. Fresh resected human PRAD specimens, including primary tumor, liver, and bone samples, were obtained through the RAP. All animal experiments were conducted in accordance with protocols approved by the University of Nebraska Medical Center Institutional Animal Care and Use Committee (IACUC; protocol # 21-048-08-FC and 16-122-11FC) and adhered to relevant institutional and national guidelines and regulations.

### Development of patient-derived xenograft and organoid xenograft models

PDX models were established by the FPBCC/UNMC RAP Patient-Derived Models Repository (RAP-PDMR) from metastatic lesions collected at rapid autopsy. Freshly procured liver and bone metastasis tissue was cut into fragments of approximately 1 mm³ and implanted subcutaneously into the flank of 8 to12 week-old male NOD scid gamma (NSG) mice (NOD.Cg-*Prkdc*^scid^*l2rg*^tm1Wjl^/SzJ; The Jackson Laboratory, Bar Harbor, ME; strain # 005557) under isoflurane anesthesia. Grafts were monitored weekly, and tumors were harvested once they reached 1 cm³. For serial passage, harvested tumors were dissected, freed of necrotic tissue, and re-implanted into recipient mice under identical conditions. The liver metastasis-derived PDX engrafted with an efficient take rate and was maintained through three serial passages. A portion of each passage was cryopreserved and fixed for histologic confirmation.

PDOX models were generated by mixing 1×10^6^ cells in a 1:1 mixture of culture medium and growth factor reduced Matrigel (Corning). Cell suspension was injected subcutaneously into 8 week-old male NSG mice (NOD.Cg-*Prkdc*^scid^*l2rg*^tm1Wjl^/SzJ; The Jackson Laboratory, Bar Harbor, ME; strain # 005557). Animals were housed under a 12h light / 12h dark cycle, and cages were changed weekly. Tumor dimensions were recorded weekly using digital calipers, and tumor volume was calculated as (length × width^2^)/2. In accordance with IACUC approved endpoints of 1cm^3^, animals were euthanized by CO_2 i_nhalation within 2 months post-engraftment. Excised tumors were divided for downstream analyses: portions were fixed for histology and processed for genomic and transcriptomic profiling, while remaining fragments were serially re-engrafted into additional NSG recipients; up to 3 serial passages were performed.

### Cell Culture and treatment

PC cell lines LNCaP, DU145, and PC3 were obtained from ATCC and RPMI-1640 medium (Gibco) supplemented with 1% penicillin/streptomycin and with 2% FBS (Corning)at 37 °C in a humidified atmosphere of 5% CO_2._

Commercial human bone marrow-derived mesenchymal stem cells (MSCs) were obtained from Lonza and ATCC. RAP-derived MSCs and healthy donor MSCs were maintained in low glucose Dulbecco’s modified Eagle’s medium (DMEM) supplemented with 10% fetal bovine serum (FBS), 1% penicillin/streptomycin, and platelet-derived growth factor BB (PDGF-BB; 10 ng/mL), and were passaged with TrypLE. For lineage differentiation, MSCs were cultured in α-Minimum Essential Medium (α-MEM) supplemented with 10% FBS, 1% penicillin/streptomycin, and lineage-specific supplements provided in the Human Mesenchymal Stem Cell Functional Identification Kit (R&D Systems).

For irradiation experiments, cells received 2 Gy of γ-irradiation and were collected 24 h post-irradiation along with non-irradiated controls. For hypoxia experiments, cells were placed in a hypoxia chamber at 2% O_2 f_or 24 h. All downstream assays were performed in a minimum of three independent biological replicates.

Cell identity was verified by short-tandem-repeat (STR) profiling. Adenocarcinoma, neuroendocrine lineage was assessed by immunoblotting and immunofluorescence for AR, NKX3.1, CK8, EpCAM, E-cadherin, synaptophysin, NCAM1, and EZH2 as detailed below.

### Patient-derived organoid culture

Patient-derived organoids (PDOs) from bone (PDO_BM1) and liver (PDO_LM1) metastases were established and maintained as previously described by [54], with minor modifications as published [52, 53, 55, 56]. Briefly, freshly procured tissue was minced into ∼1 mm^3^ fragments and digested in advanced DMEM/F12 (adDMEM/F12; Thermo Fisher Scientific) containing 5 mg/mL collagenase type II (Thermo Fisher Scientific), 10 µM Y-27632 (Tocris), and primocin (InvivoGen; 1X) for 1-2 h with continuous rocking in an orbital shaker at 37 °C. The digest was further dissociated by brief incubation in TrypLE Express (Thermo Fisher Scientific) at 37 °C for 5-10 min, neutralized in adDMEM/F12 supplemented with 10% FBS, filtered through a 70 µm cell strainer, and pelleted at 300 x g for 5 min. Cell pellets were resuspended in growth factor reduced Matrigel (Corning) and plated as 30-50 µL droplets in pre-warmed 24 well plates. After polymerization at 37 °C for 15-20 min, droplets were overlaid with complete human prostate organoid medium consisting of adDMEM/F12 supplemented with 1X GlutaMAX, 10 mM HEPES, 100 U/mL penicillin and 100 µg/mL streptomycin, 1X B27 (Thermo Fisher Scientific), 1.25 mM N-acetyl-L-cysteine, 10 mM nicotinamide, 5 ng/mL recombinant human EGF (PeproTech), 500 ng/mL recombinant human R-spondin 1 (R&D Systems), 100 ng/mL recombinant human Noggin (PeproTech), 500 nM A83-01 (Tocris), 10 ng/mL recombinant human FGF10 (PeproTech), 5 ng/mL recombinant human FGF2 (PeproTech), 1 µM prostaglandin E2 (Tocris), 10 µM SB202190 (Sigma-Aldrich), and 1 nM dihydrotestosterone (Sigma-Aldrich). Y-27632 (10 µM) was included for the first 2-3 days after seeding and after each passage to support single-cell viability. Media was refreshed every 2-3 days. Organoids were passaged every 5 days by mechanical disruption with a P1000 pipette followed by brief TrypLE digestion (5-10 min at 37 °C), and were re-embedded in fresh Matrigel at a 1:3 split ratio. Cultures were maintained at 37 °C in a humidified atmosphere of 5% CO₂. The bone and liver metastasis-derived cell lines were cultured in the same media as described above and maintained at 37°C and 5% CO_2._

To model the bone marrow niche, we established a co-culture system pairing the bone metastasis-derived PDO (PDO_BMO) with an iPSC-derived bone marrow organoid. Bone marrow organoids were generated by directed differentiation of iPSCs through mesenchymal, endothelial, and hematopoietic lineages following published protocols [94, 95, 98]. Mature organoids were selected for co-culture at day 18 of differentiation, when vascular network formation was confirmed. PDO_BMO were dissociated into single cells and seeded onto the iPSC-derived bone marrow organoid in organoid culture medium. Co-cultures were maintained in organoid culture medium at 37 °C and 5% CO₂ for the duration, with medium refreshed every 3 days. Co-culture was assessed by immunofluorescence.

### Isolation of MSCs from PC rapid autopsy patient bone chips

Bone fragments (∼1 mm^3^) extracted from patient spinal metastases were rinsed in 1X PBS and digested in advanced DMEM/F12 (adDMEM/F12; Thermo Fisher Scientific) containing 5 mg/mL collagenase type II (Thermo Fisher Scientific), 10 µM Y-27632 (Tocris), and primocin (InvivoGen; 1X) for 1-2 h with continuous rocking in an orbital shaker at 37 °C. The digest was further dissociated by brief incubation in TrypLE Express (Thermo Fisher Scientific) at 37 °C for 5-10 min, neutralized in adDMEM/F12 supplemented with 10% FBS, filtered through a 70 µm cell strainer, and pelleted at 300 x g for 5 min. The supernatant and cell pellet were then plated in 12-well culture plates for cell expansion.

## METHOD DETAILS

### Adipocyte Differentiation Assay

MSCs were seeded in a 24 well plate and cultured until confluent. Cells were then cultured in alpha-MEM with adipogenic supplement (R&D Systems) for 8-12 days with fresh media replacement every 3-4 days. At the end of the treatment period, cells were washed with 1X PBS and fixed in 10% neutral buffered formalin for 1 hour. Cells were washed twice with PBS, once with 60% isopropanol, and then allowed to air dry in the well plate for 10 minutes. Adipocytes were identified by staining of lipid content using Oil Red O (Sigma-Aldrich) for 10 minutes and then washed twice with deionized water. Representative images per well were taken under brightfield microscopy at 20X magnification. Differentiation assays were performed a minimum of 3 times.

### Osteoblast Differentiation Assay

MSCs were seeded in a 24-well plate and cultured until confluent. Cells were then cultured in alpha-MEM medium with osteogenic supplement (R&D Systems) for 12-21 days with fresh media replacement every 3-4 days. At the end of the treatment period, cells were washed once with 1X PBS and fixed in 10% neutral buffered formalin for 15 minutes. To determine the amount of osteogenesis, cell-derived calcium deposits were stained with Alizarin Red solution. Cells were washed twice with deionized water and stained with 2% Alizarin Red solution (LabChem), pH 4.3, for 45 minutes in the dark, then washed three times with deionized water and the well plate allowed to air dry overnight. To extract and quantify the stained mineral, 10% acetic acid was added to each well and incubated at room temperature for 30 minutes. Each well was scraped with a pipette tip to collect cells from the plate; cells and supernatant were then transferred to microcentrifuge tubes and heated at 85 degrees Celsius for 10 minutes. The reaction was stopped by placing tubes on ice for 5 minutes and then centrifuged for 15 minutes at 20,000 x g. 125 uL of supernatant was transferred to a new tube to which 50 uL of 10% ammonium hydroxide was added. Tubes were mixed, and 50 uL samples were placed in a clear flat-bottomed 96-well plate. Absorbance was measured at 405 nm.

### Flow Cytometry

MSC cell surface markers were measured using Human Mesenchymal Stem Cell Multi-Color Flow Kit (R&D Systems) according to the manufacturer’s protocol.

### Gene Expression

RNA was isolated from MSCs in complete growth media using Trizol reagent, according to the manufacturer’s protocol. RNA (1 μg) was used to synthesize cDNA using qSCRIPT Super mix (Quantabio). Real-time quantitative PCR (RT-qPCR) was performed using Perfecta SYBR Green FastMix (Quantabio) and run using Bio-Rad CFX Real-Time System. PCR conditions were as follows for all primer sequences (Table 1): Step 1: 95° 30 s; Step 2: 95° 5 s, 57° 15 s, 72° 10 s, 95° 10 s (× 39 cycles); Step 3: Melt curve 65° 95° increments of 0.5° for 5 s. Relative gene expression was calculated using the ΔΔCT method, normalized to expression of housekeeping gene 36B4.

### Immunoblotting

Cells were lysed in freshly prepared RIPA buffer supplemented with 1% protease/ phosphatase inhibitor cocktail (Sigma) and 1% EDTA (Sigma). Protein concentration was determined using the Bio-Rad DC Protein Assay Kit, with absorbance read on a BioTek Synergy H1 plate reader. Equal amounts of protein (15-20 µg) were resolved on 12% Bis-Tris gels (Invitrogen) at 100 V and transferred to Immobilon-P PVDF membranes. Membranes were blocked in 5% non-fat milk in TBST for 1 h at room temperature and incubated overnight at 4°C with primary antibodies at an appropriate dilution in 5% milk. Membranes were then washed and incubated for 1 h at room temperature with HRP-conjugated secondary antibody at 1:3,000 in 5% milk. Signal was detected using SignalFire TM ECL reagent (Cell Signaling Technology) and imaged on the ChemiDoc imaging system (Bio-Rad). The same membranes were re-probed with an antibody against beta-actin as a loading control. A complete list of antibodies, including catalog numbers and RRIDs, is provided in Table S1.

### Reactive oxygen species (ROS) detection

Following irradiation or hypoxia exposure as described above, live cells were incubated with dihydroethidium (DHE; Thermo Fisher Scientific, MA) (5 µM) or 5-(and-6)-chloromethyl-2′,7′-dichlorodihydrofluorescein diacetate (CM-H_2D_CFDA; Thermo Fisher Scientific, MA) (5 µM) for 30 min at 37 °C. Fluorescence images were acquired on a Zeiss LSM 800 confocal microscope at the UNMC Advanced Microscopy Core Facility.

### Histology

Patient-derived organoids (PDOs) were fixed in 4% paraformaldehyde (Electron Microscopy Sciences, Hatfield, PA) for 30 min at room temperature, washed with PBS, and resuspended in 2% low-melting-point agarose in PBS before processing. Patient bone metastasis specimens were collected at rapid autopsy and fixed in 10% neutral-buffered formalin for 24 h at room temperature. Fixed specimens were decalcified in 0.5 M EDTA, pH 7.4, and formic acid based decalcifier for 7 days at 4C, with the solution refreshed every 3 days. Decalcified tissue was rinsed in PBS, dehydrated through a graded ethanol series, cleared in xylene, and embedded in paraffin. Sections were cut at 5 µm for downstream hematoxylin and eosin staining. Patient-derived xenograft (PDX) and patient-derived organoid xenograft (PDOX) tissues were fixed in 4% paraformaldehyde overnight at room temperature and washed with PBS. All samples were transferred to 70% ethanol and submitted to the UNMC Tissue Sciences Facility (TSF) for paraffin embedding and sectioning. Every 10th or 20th 5-µm section was stained with hematoxylin and eosin (H&E) using a Tissue-Tek Prisma Plus automated stainer (Sakura Finetek USA, Torrance, CA). Slides were scanned at 40× magnification on an Aperio CS2 scanner using Aperio software (Leica Biosystems, Wetzlar, Germany).

Immunofluorescence staining was performed as previously described [52, 113], with minor modifications. Sections from PDOs, PDXs, PDOXs, and patient liver and bone metastases were deparaffinized with HistoChoice Clearing Agent (VWR). Antigen retrieval was performed by microwave heating for 15 min in one of the following buffers, selected according to the antibody: 10 mM citrate buffer (pH 6.0), 1 mM EDTA (pH 8.0), or Tris-EDTA buffer (10 mM Tris, 1 mM EDTA, 0.01% Tween 20, pH 9.0). Sections were quenched in 50 mM glycine in PBS for 15 min at room temperature, then blocked in PBS containing 10% normal goat serum and 0.3% Triton X-100 for 1 h at room temperature. Primary antibodies (Table S2) were applied for the durations indicated (overnight to 48 h), after which sections were washed three times in PBS (5 min each) and incubated with Alexa Fluor 488, 555, or 657 donkey anti-rabbit secondary antibody (1:250; Thermo Fisher Scientific) for 1.5 h at room temperature. Nuclei were counterstained with 1 mg/mL 4′,6-diamidino-2-phenylindole (DAPI) for 5 min at room temperature. Sections were rinsed in distilled water and mounted with ProLong Gold Antifade Mountant (Thermo Fisher Scientific). Confocal images were acquired on a Cytation C10 Confocal Imaging Reader (Agilent) and Zeiss LSM710 confocal microscope (Carl Zeiss Microscopy) at the UNMC Advanced Microscopy Core Facility (AMCF). Images were analyzed using the respective software: Aperio Digital Pathology Imaging System (Leica) QuPath, Gen5 software (Agilent), and Zen software (Carl Zeiss Microscopy). Whole-slide scans were obtained using a Zeiss Cell Discoverer 7 or Zeiss Axio Scan with Zen software at the UNMC Advanced Microscopy Core Facility.

Whole-mount immunofluorescence of organoids was performed as previously described [52, 113]. Organoids were washed in PBS and fixed in 4% paraformaldehyde in PBS overnight at 4 °C. Fixed organoids were blocked in PBS containing 1% Triton X-100 and 1% FBS for 60 min at room temperature, then incubated for 1 h at room temperature each with primary and secondary antibodies diluted in PBS containing 0.5% Triton X-100 and 10% normal goat serum. Nuclei were counterstained with DAPI. Z-stacks were acquired on a Zeiss LSM710 confocal microscope (Carl Zeiss Microscopy).

### RNA sequencing and analysis

RNA sequencing of patient-derived organoids was performed using Illumina’s NovaSeq platform with unstranded library preparation (Azenta). Paired-end reads were generated with > 19 million reads per sample and mean quality score > 35. The transcript data in fastq files were aligned to the genome using STAR (version 2.7.9a); a genome index was generated using the GRCh 38 primary assembly genome fasta. QC was conducted prior to and after alignment using FastQC and PicardTools, respectively. Transcript quantification was obtained using either STAR counts or salmon (version 1.5.2). Differential analysis was conducted using DESeq2, and fragments per kilobase of transcript per million mapped reads (FPKM) were exported.

### Patient disease classification

To classify the transcriptomes of our patient-derived organoids, we identified 691 PC patients with open access RNA sequencing data on the Genomic Data Commons (GDC) portal. The samples were derived from 3 different studies and included normal tissue from the prostate and tumors from the primary site and metastatic site. As a result of its harmonized bioinformatics pipeline, the GDC hosts raw gene expression count files following alignment of RNA-seq transcripts to the genome using STAR. The unstranded counts from these files were read into R along with unstranded counts from the organoids (derived by similar genomic alignment using STAR). The resulting raw count matrix was normalized using DESeq2 for principal component analysis. Batch correction for the study of origin was applied using limma’s removeBatchEffect method. To achieve more granular classification, we obtained NEPC and adenocarcinoma scores based on the classifier created by Beltran et al 2016. FPKM values were collected for the reference set of 70 genes (additional genes: *PTEN*, *TP53*) from the RNA-seq of PDOs and PC cell lines in the dataset from Smith et al. [136]. The Pearson’s correlation coefficient was determined between the PDOs/cell lines and the NEPC reference vector and reported as the NEPC score. Samples with an NEPC score ≥ 0.40 are classified as putative CRPC-NE tumors.

Raw WES and RNA-seq FASTQs were downloaded from the NCI PDMR database for the independently identified RAP patient sample (Patient ID:465922, samples 331-R2 and 331-R4; https://dctd.cancer.gov/drug-discovery-development/reagents-materials/pdmr). These included sequencing results from the original tumor (ORIGINATOR) and PDX tumors. Raw FASTQ files were binned into human or mouse reads using relevant reference genomes (GRCh38, GRCm39) in bbsplit. WES fastqs were aligned to GRCh38 reference using bwakit. Somatic variants were called using the Genome Analysis Toolkit “Mutect2” in tumor-only mode with the following parameters: germline-resource (hg38.af.only.gnomad.vcf.gz) and af-of-alleles-not-in-resource 5e-8; calls were then filtered using FilterMutectCalls with default parameters. PureCN was used to determine sample purity and ploidy and assess loss of heterozygosity and gene-level copy number. Purity-and ploidy-corrected segment-level copy number was then obtained using cnvkit call, and copy-number plots were produced with the CopyNumberPlots R package. RNA-seq fastqs were processed as described previously [137, 138]. VST-normalized counts and FPKM normalized counts were obtained to facilitate comparison to organoid samples and to generate NEPC scores, respectively.

### Quantification and statistical analysis

Statistical analyses were performed using GraphPad Prism (v 11; GraphPad Software, San Diego, CA). Data are presented as mean ± SD or mean ± SEM of at least three independent biological replicates unless otherwise indicated. Comparisons between two groups were performed using two-tailed Student’s t-test or Mann-Whitney U test; comparisons among three or more groups were performed using one-way ANOVA followed by Tukey’s/Dunnett’s post hoc test. A p-value < 0.05 was considered statistically significant. Specific statistical tests, sample sizes (*n*), and p-values for each experiment are reported in the corresponding figure legends.

## FIGURE LEGENDS

**Supplementary Table 1.** Tables representing PDMR samples (sheet1) and the OncoKB^TM^ analysis (sheet2) list of gene mutations identified through whole exome sequencing of RAP liver metastasis tissue.

**Supplementary Figure 1.** Immunofluorescence profiling of PC RAP liver metastasis tissue. (A) Whole tissue immunofluorescence staining of the liver metastasis section procured via the PC RAP for six markers: KRT8/18 (luminal epithelial marker), SYP (synaptophysin, neuroendocrine marker), NCAM1 (neuroendocrine adhesion molecule), EZH2 (epigenetic modifier), Ki-67 (proliferation marker), and NKX3.1 (AR-regulated luminal transcription factor). For each marker, DAPI nuclear counterstain (blue, left panel), marker channel (green, middle panel), and merged image (right panel) are shown. Scale bars, 1 mm.

**Supplementary Figure 2.** High-magnification immunofluorescence of PC RAP liver metastasis tissue. (A) Immunofluorescence staining of liver metastasis tissue: SYP (synaptophysin, neuroendocrine marker), EZH2 (epigenetic modifier), and NKX3.1 (AR-regulated luminal transcription factor). For each marker, DAPI nuclear counterstain (blue, left panel), marker channel (green, middle panel), and merged image (right panel) are shown. Scale bars, 100 µm.

**Supplementary Figure 3.** Co-expression analysis of luminal, neuroendocrine, proliferation, epithelial, and mesenchymal markers within the same liver metastasis. (A) Dual color immunofluorescence of liver metastasis sections from PC-RAP tissue, stained for DAPI (blue) and the following marker pairs: KRT8/18 (green) and NCAM1 (red); KRT8/18 (green) and EZH2 (magenta); Ki-67 (green) and E-cadherin (CDH1; magenta); and Ki-67 (green) and vimentin (VIM; magenta). Individual DAPI, and marker 1, and marker 2 channels are shown in grayscale, with the corresponding merged image at right. Boxed regions in the merged panels are enlarged in the inset to highlight cellular co-localization. Scale bars, 100 μm.

**Supplementary Figure 4.** Superoxide (DHE) levels in PC RAP-derived cell lines under radiation and hypoxia. (A) Representative fluorescence microscopy images of bone and (B) liver metastasis-derived cells stained with DHE (superoxide, red) under control and 2 Gy radiation conditions. Scale bars, 20 µm.

**Supplementary Figure 5.** Patient-derived xenografts and spontaneous lymph node metastases retain the luminal and neuroendocrine marker profile of the parental liver metastasis tissue. (A) Immunofluorescence of a patient-derived xenograft (PDX) established from the PC-RAP liver metastasis tissue, stained for DAPI (blue) and either SYP or EZH2 (green). DAPI and target channels are shown in greyscale alongside the merged image. (B) Immunofluorescence of a lymph node metastasis arising from the liver-metastasis derived PDX in (A), stained for DAPI (blue) and the indicated markers (green): KRT8/18, NCAM1, and Ki-67. Scale bars, 100 μm.

**Supplementary Figure 6.** A KRT8/18⁺ / KRT5/14⁻ luminal cytokeratin profile is conserved across PDX, PDOX, and lymph node metastases. Co-expression immunofluorescence analysis of DAPI (blue), basal cytokeratin KRT5/14 (red), and luminal cytokeratin KRT8/18 (green) across patient-derived models established from the PC-RAP liver metastasis: (A) parental liver-metastasis PDX; (B&C) patient-derived orthotopic xenograft (PDOX: LMO, BMO)and (D) a lymph node metastasis arising from the BMO PDOX in (C). For each model, individual DAPI, KRT5/14, and KRT8/18 channels are shown in greyscale alongside the merged image. Boxed regions in the merged panels are enlarged in the inset. All models retain a predominantly KRT8/18⁺ / KRT5/14⁻ luminal cytokeratin profile, with negligible basal cytokeratin expression. Scale bars, 100 μm.

## ABBREVIATIONS

ADT: androgen deprivation therapy
CRPC: castration-resistant prostate cancer
LN: lymph node
mCRPC: metastatic castration-resistant prostate cancer
NEPC: neuroendocrine prostate cancer.
DMSO: dimethyl sulfoxide
FFPE: formalin-fixed paraffin-embedded
PDO: patient-derived organoid
PDOX: patient-derived organoid xenograft
PDX: patient-derived xenograft
LOH: loss of heterozygosity
PDMR: Patient-Derived Models Repository
PTEN: phosphatase and tensin homolog
RB1: retinoblastoma 1
WES: whole exome sequencing
CK: cytokeratin
DAPI: 4′,6-diamidino-2-phenylindole
H&E: hematoxylin and eosin
NCAM1: neural cell adhesion molecule 1
PDO_BMO: bone metastasis-derived patient organoid
PDO_LMO: liver metastasis-derived patient organoid.
FPKM: fragments per kilobase of transcript per million mapped reads.
DAPI: 4′,6-diamidino-2-phenylindole
ROS: reactive oxygen species
CRPC: castration-resistant prostate cancer
DAPI: 4′,6-diamidino-2-phenylindole
H&E: hematoxylin and eosin
iPSC: induced pluripotent stem cell.

## Notes

### Competing Interest Statement

The authors have declared no competing interest.

