## Supplemental figures for "An Integrated Preclinical Platform for Lethal Neuroendocrine Prostate Cancer from Rapid Autopsy Bone and Liver Metastases"

**A** <u>Immunostaining: Liver Metastasis</u>

**Supplementary Figure 1**

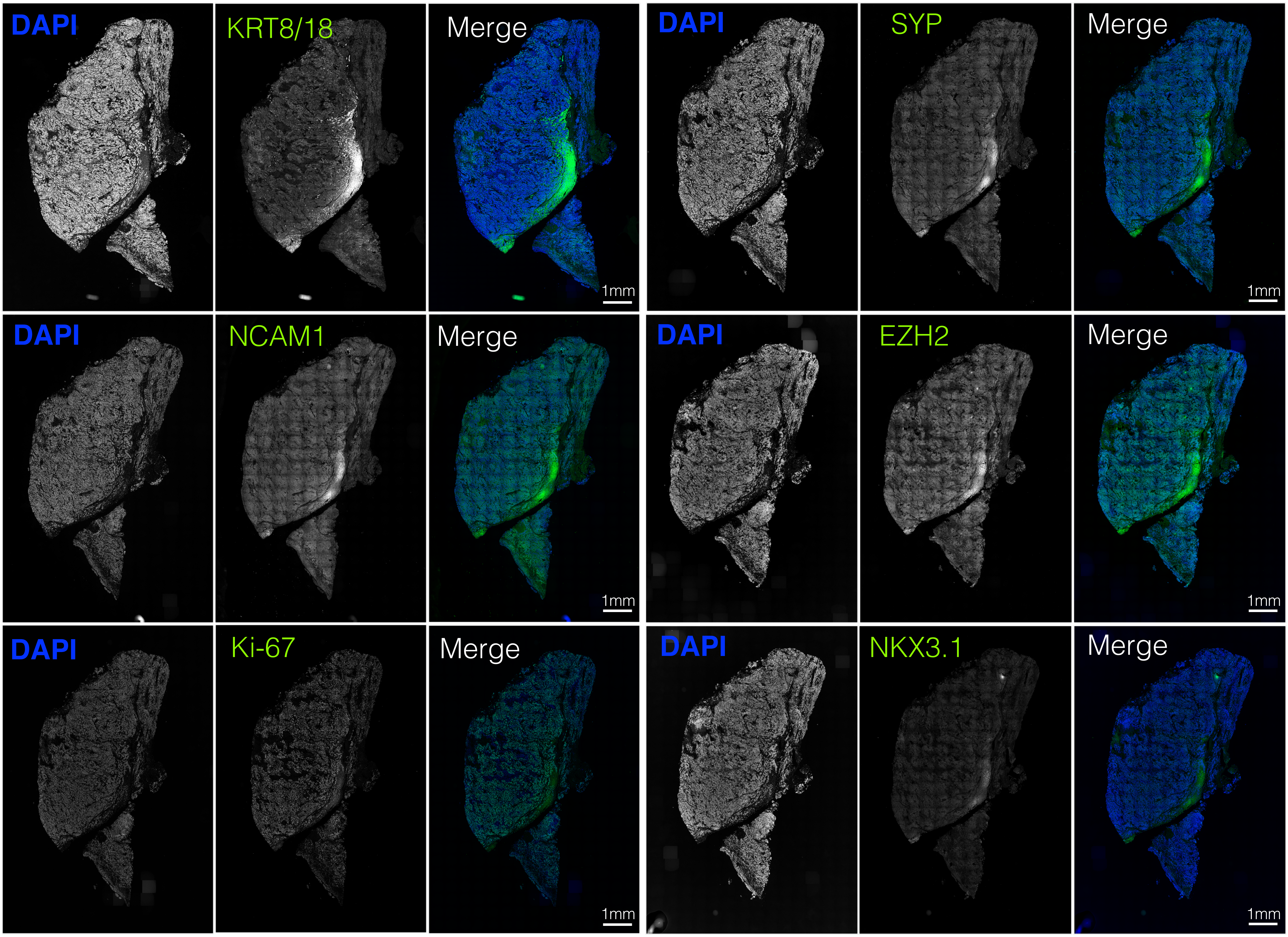

Supplementary Figure 2

**A** Immunostaining: Liver Metastasis

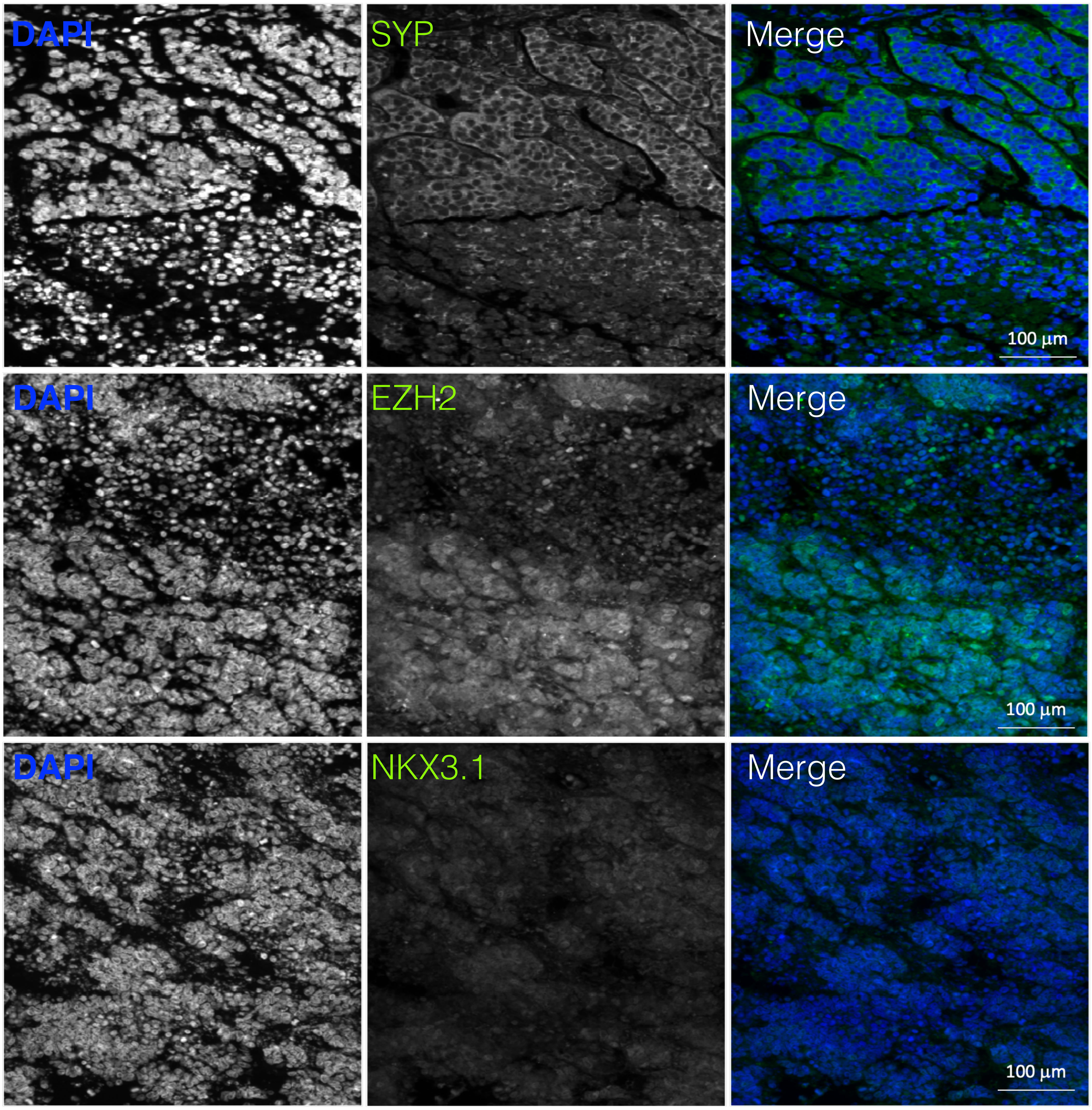

Supplementary Figure 3

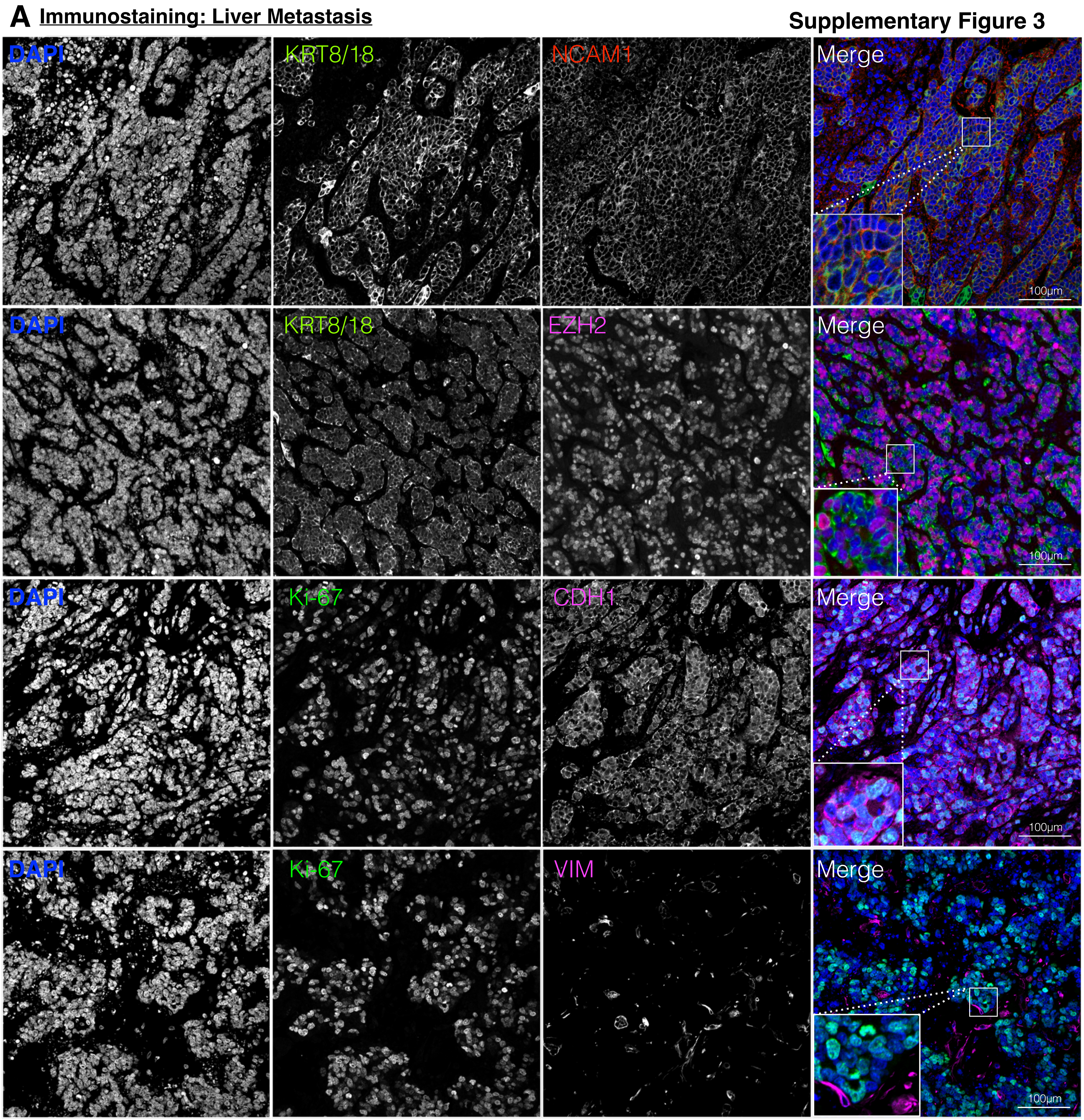

### Supplementary Figure 4

**A** Bone metastasis derived cell line

**B** Liver metastasis derived cell line

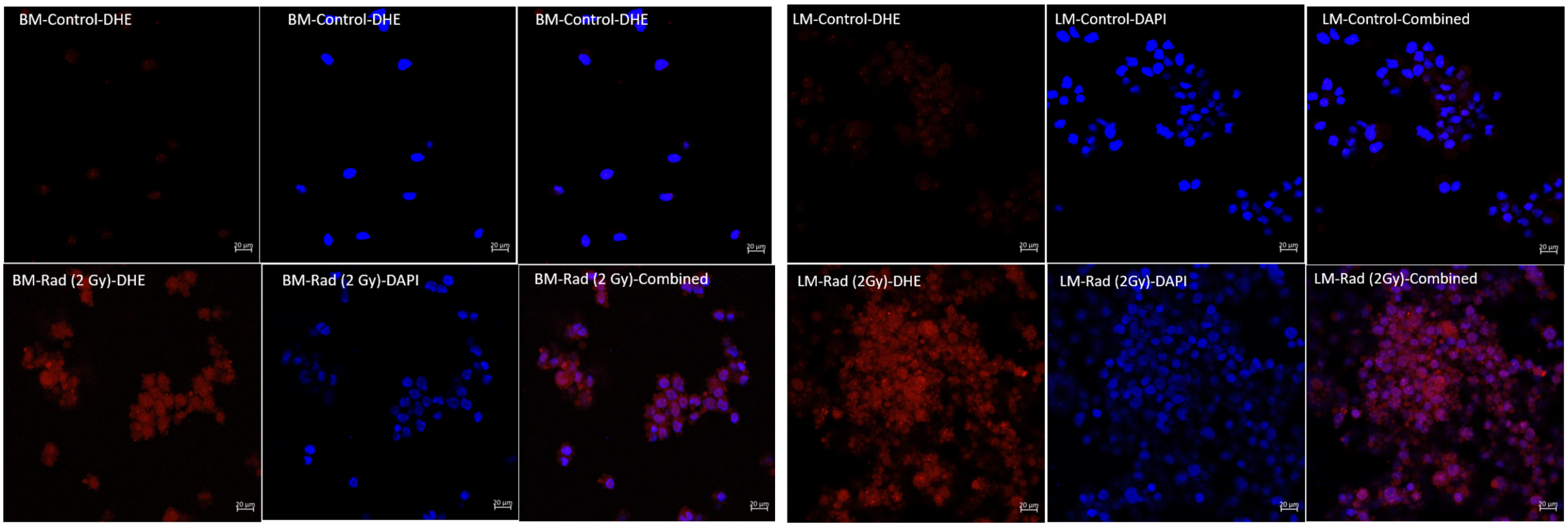

**Supplementary Figure 5**

**A** **Immunostaining: PDX (Liver metastasis derived)**

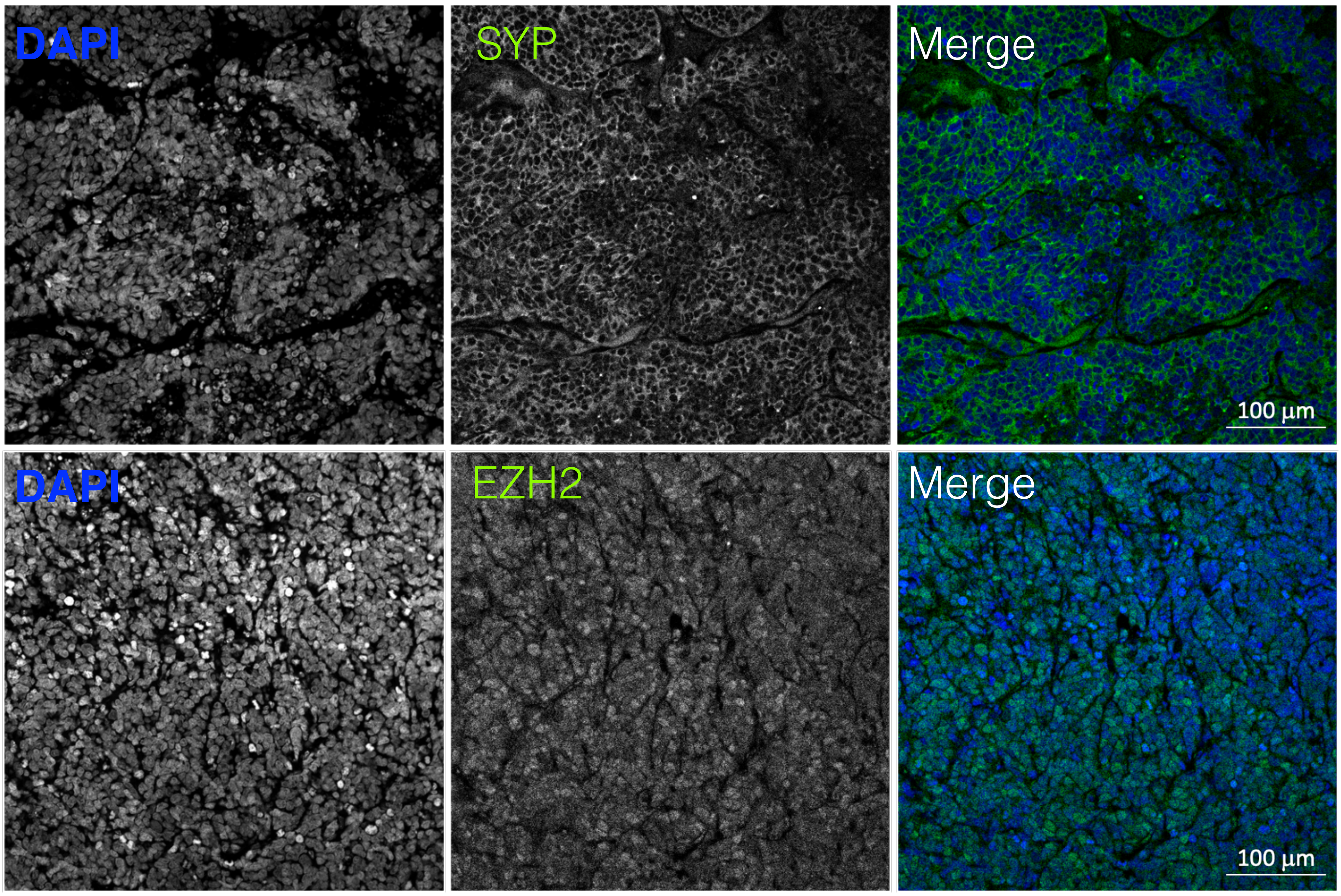


**B** **Immunostaining: Lymph node metastasis from PDX (Liver metastasis)**

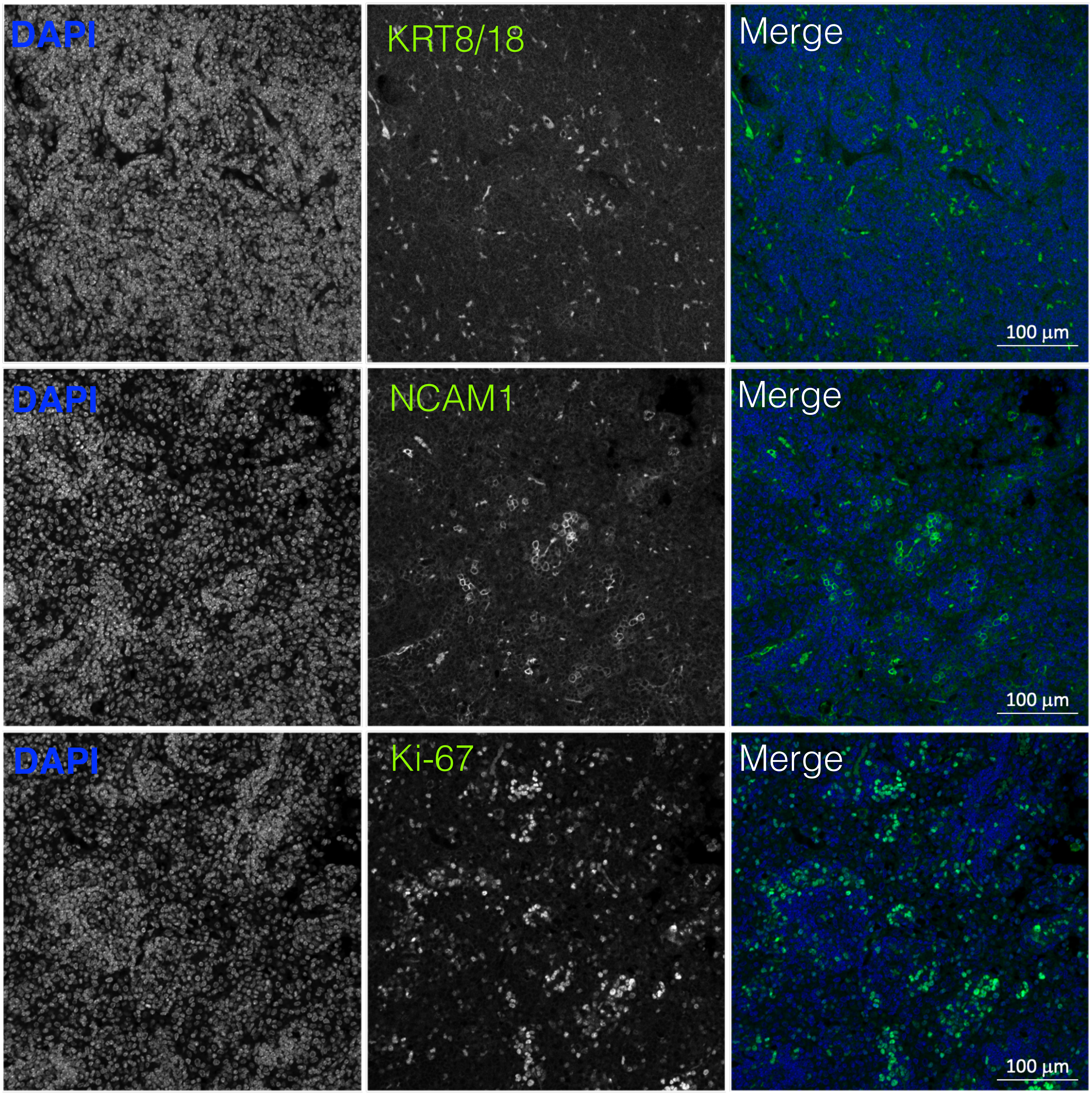

**Supplementary Figure 6**

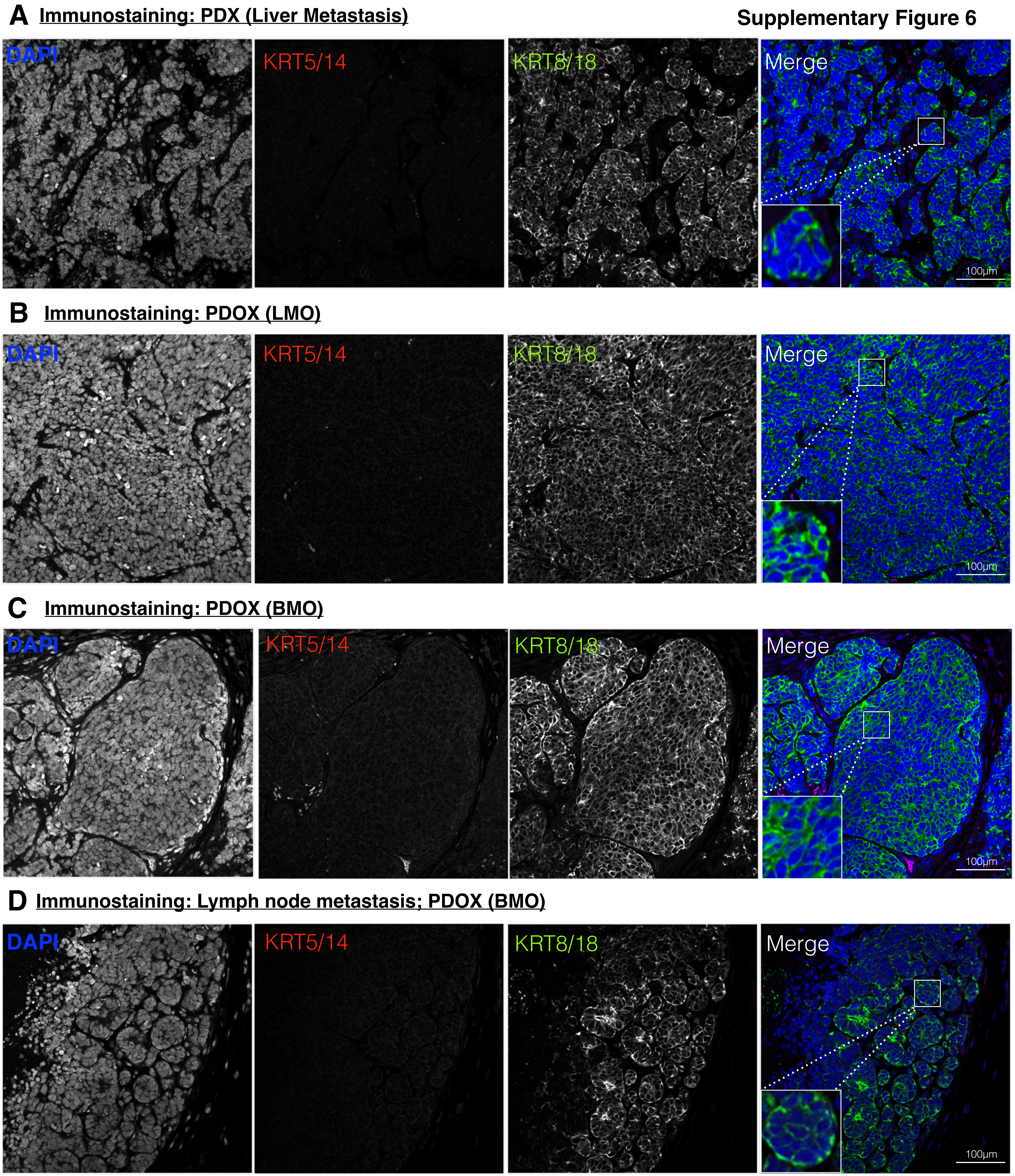
